# Perturbing proteomes at single residue resolution using base editing

**DOI:** 10.1101/677203

**Authors:** Philippe C Després, Alexandre K Dubé, Motoaki Seki, Nozomu Yachie, Christian R Landry

## Abstract

Base editors derived from CRISPR-Cas9 systems and DNA editing enzymes offer an unprecedented opportunity for the precise modification of genes, but have yet to be used at a genome-scale throughput. Here, we test the ability of an editor based on a cytidine deaminase, the Target-AID base editor, to systematically modify genes genome-wide using the set of yeast essential genes. We tested the effect of mutating around 17,000 individual sites in parallel across more than 1,500 genes in a single experiment. We identified over 1,100 sites at which mutations have a significant impact on fitness. Using previously determined and preferred Target-AID mutational outcomes, we predicted the protein variants caused by each of these gRNAs. We found that gRNAs with significant effects on fitness are enriched in variants predicted to be deleterious by independent methods based on site conservation and predicted protein destabilization. Finally, we identify key features to design effective gRNAs in the context of base editing. Our results show that base editing is a powerful tool to identify key amino acid residues at the scale of proteomes.

## Introduction

Recent technical advances have allowed the investigation of the genotype-phenotype map at high resolution by experimentally measuring the effect of all possible nucleotide substitutions in a short DNA sequence. While saturated mutagenesis informs us on the effect of many mutations, it usually covers a single locus or a fraction of it^1, 2^. Because such data is only available at sufficient coverage for a very small number of proteins, general rules on substitution effects must be extrapolated to other, often unrelated proteins. At a lower level of resolution, genome-scale mutational data has mostly been acquired through large-scale loss-of-function strain collections, where the same genetic change (for example, complete gene deletion) is applied to all genes^3–5^. This approach is a powerful way to isolate each gene’s contribution to a phenotype, including fitness, but limits our understanding of the role of specific positions within a locus.

CRISPR-Cas9 based approaches usually cause protein loss of function through indel formation^6^ or by modifying gene expression levels^7–9^ at many loci in parallel. Again, these approaches generally limit the information gain to one perturbation per locus. There is therefore a strong tradeoff between the resolution of the existing assays and the number of loci or genes investigated. Recent developments in the field now allow for the exploration of the effects of many mutations per gene across the genome. For instance, in yeast, methods for high throughput strain library construction have allowed the measurement of thousands of variant fitness effects in parallel across the genome^10–14^. These approaches rely on CRISPR-Cas9 based genome modifications requiring the formation of double-strand breaks followed by repair using donor DNA, which often depends on complex strain and plasmid constructions. An alternative approach would be to use base editors, which allow the introduction of the mutations of interest directly in the genome by direct modification of DNA bases rather than DNA segment replacement.

Base editors use DNA modifying enzymes fused to modified Cas9 or Cas12 proteins to create specific point mutations in a target genome^15–17^. Such base editors have recently been used to perform site-specific forward mutagenesis in human cell lines. The two main approaches, Targeted AID-mediated mutagenesis (TAM)^18^ and CRISPR-X^19^, target specific regions of the genome where they induce mutations randomly. This generates a library of mutant genotypes that can be competed to find beneficial and deleterious variants under selective pressure. As the relative fitness measurements depend on targeted sequencing of the locus of interest, these approaches are difficult to adapt to high throughput multiplexed screens where tens of thousands of sites can be targeted within the same gRNA libraries.

Here, we present a method that bridges the flexibility of Target-AID mutagenesis and the multiplexing capacities of genome editing depletion screens. By using a base editor with a narrow and well-defined activity window^15^, we selected gRNAs generating a limited number of predictable edits in yeast essential genes. This allowed us to use gRNAs as a readout for the effect of the mutations, similar to commonly used barcode-sequencing approaches to measure fitness effects.

## Results

### Design of a base editing library targeting essential genes

We used Target-AID mutagenesis to simultaneously assess mutational effects at over 17,000 putative sites in the yeast genome. We scanned yeast essential genes for sites amenable to editing by the Target-AID base editor as well as targets with other specific properties, including intronic sequences. Because all essential genes have the same qualitative fitness effects when deleted^20^, focusing on these genes allowed us to limit the variation in fitness that could be due to the relative importance of individual genes for growth rather than to the importance of specific positions within a locus. We excluded gRNAs that did not target between the 0.5th and 75th percentile of the length of annotated genes to limit position biases that could influence the efficiency of stop-codon generating guides^21, 22^.

To associate each gRNA in the library to specific base editing outcomes, we developed a simple model based on the yeast data included in the original Target-AID manuscript as well as our own work^15, 23^. First, we expected that editing would mostly result in genotypes where only one nucleotide is edited in the activity window of the editor. Second, we predicted that the editing outcomes would mainly consist of C to G and C to T mutations and that the abundance of C to A products will be negligible. Finally, we expected that editing frequency ranks would follow the editing activity rankings already known from the initial characterization of Target-AID. Based on these criteria, we filtered out potential target sites where all three high editing rate positions (−19,−18 and −17) or those where both position −18 and −17 are cytosines and kept the remaining sites for inclusion in the gRNA library. The resulting library contained 40 000 gRNAs, of which ∼35 000 targeted essential gene coding sequences and ∼5000 other target types as shown in Supplementary Figure 1.

Over 75% of target sequences in this set contained only one or two Cs in the extended activity window (positions −20 to −14), and as expected a general enrichment for cytosines in the high activity window (Supplementary Figure 2A-B). Because the goal of our experiment was to link specific mutations to fitness effects, co-editing of multiple nucleotides using an editor which does not channel mutations to a specific outcome has the potential to obscure the genotype responsible for a fitness effect. To take this into account, we placed each gRNA in a co-editing risk category based on the presence and positions of cytosines in the activity window (See methods). Based on this metric, we found that over 80% of gRNAs fell either in the very low or low risk category (Supplementary Figure 2C). If co-editing occurs, but the other mutated cytosine is part of the same codon as the intended target site, then any resulting fitness effects can still be linked to the perturbation of a specific amino acid. We found the proportion of gRNAs in the library for which this is true to be over 50%: when co-editing risk category is taken into account, the proportion reaches ∼90% (Supplementary Figure 2D). As Target-AID is known to perform processive editing, a high co-editing risk might also be linked to higher overall editing rate^15^.

### Measurement of mutagenesis rate and outcomes of library gRNAs

While the repair product outcomes of edits for gRNAs can be predicted with varying levels of accuracy for CRISPR-Cas9-based editing^24^, no such tools are available yet for base editing applications. As such, the model we used to associate gRNAs in our library to mutational outcomes is only a parsimonious deduction based on the original Target-AID data and our previous work^15, 23^. Furthermore, evaluating the activity of gRNAs for base editing remains difficult^25^. The measurement of fitness effects is not associated with a direct simultaneous measurement of mutagenesis rate in our experiment. As such, the absence of fitness effects for a gRNA can both be explained by either non-functional or low editing, or successful editing that resulted in mutations with no detectable fitness effects^23^. As our experiment focuses on the impact of targeted mutations on cell growth, the first group can be seen as false negatives, and the second as true negatives. While we can modulate the gRNA abundance variation threshold to minimize the risk of false positives, additional experimental data on mutagenesis success rates and editing outcomes was required to assess which type of negative results would be dominant in our experiment.

To evaluate the performance of our model and the functionality of the library gRNAs, we performed a base editing time course experiment where mutagenesis rates and outcomes were measured by deep sequencing of the edited genomic loci (Supplementary Figure 3). To gain insights on the mutagenesis outcomes of different editing scenarios, we selected guides with different predicted patterns of cytosine presence in the Target-AID activity window (Figure 1A). We included 9 guides from the library isolated from the library quality control process (see methods), as well as three control gRNAs respectively targeting the pseudogene YCL074W, the non-essential gene *VPS17*, and *ADE1*, which can be used as a phenotypic marker. Most gRNAs could efficiently edit their respective targets, with 9 out 12 gRNAs reaching mutation rates of 50% or higher (Figure 1B), consistent with previous results^15, 23^. Replicates were highly correlated along different measurements with editing rates at the *CAN1* co-editing site being highly consistent (Supplementary Figure 4A-E). Only the gRNA targeting *SES1* was found to be inactive, and as such was excluded from downstream analysis. The very low editing rate observed for the gRNA targeting *SES1* is an example of unknown factors affecting mutagenesis efficiency that leads to false negatives in large-scale experiments.

**Figure 1.**
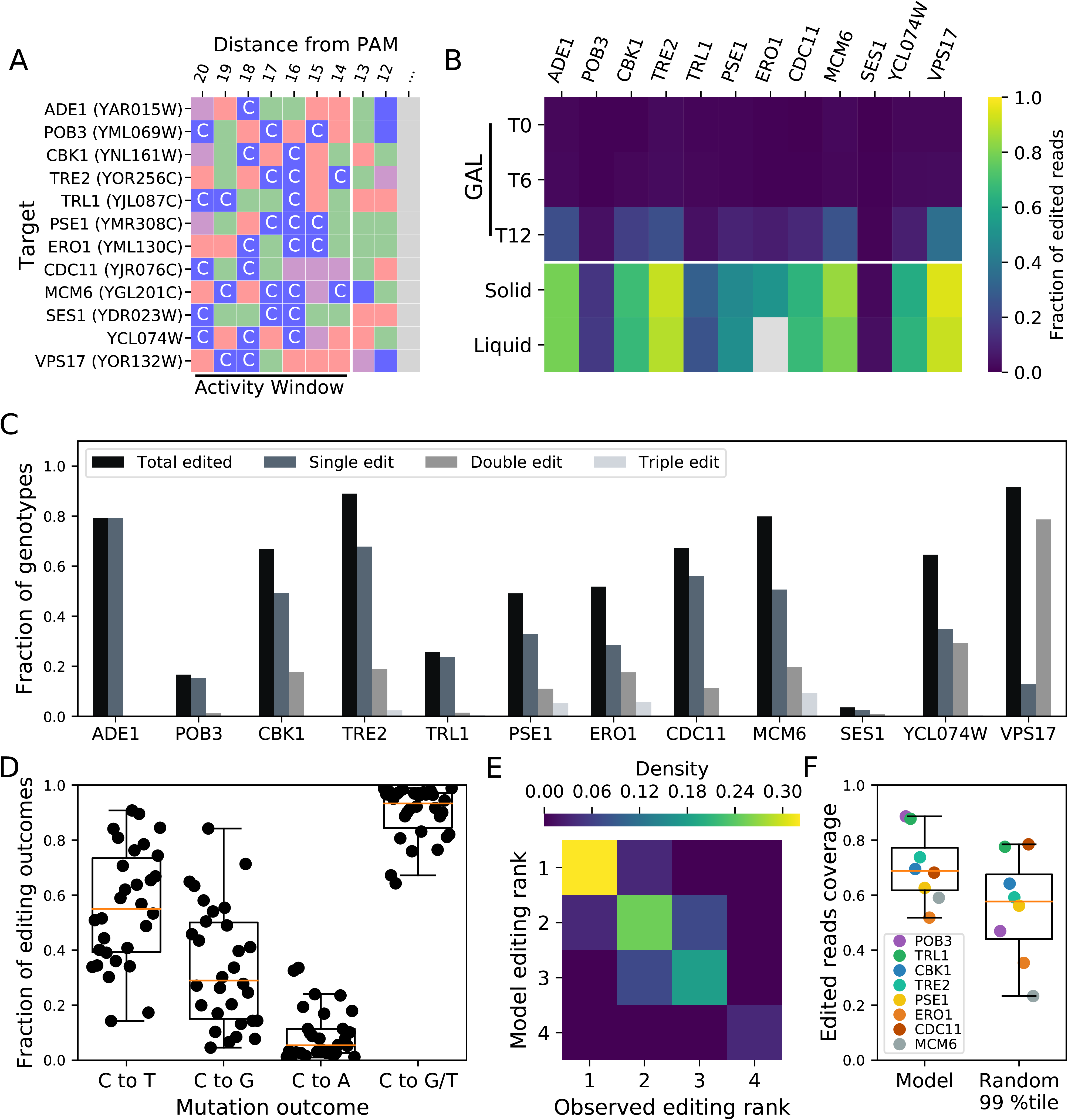
A simple parsimonious model predicts the most probable outcomes of Target-AID mutagenesis. **A)** gRNAs included in the time course base editing experiment had diverse C content profiles in the Target-AID activity window. Nucleotides are color coded: guanines are purple, thymines are red, adenines are green and cytosines are blue. **B)** Overall fraction of edited reads for all target sites rate along timepoints in the experiment: T0 (start of induction), T6 (mid induction), T12 (end of induction). The solid time point represents surviving cells plated after galactose induction, while the liquid time point represents the cell population after canavanine co-selection. Amplification of the *ERO1* target site from the liquid recovery time points was unsuccessful (shown in grey), and as such the solid recovery time point was used instead for the other analysis steps. **C)** Fraction of genotypes with different numbers of edited nucleotides in the Target-AID activity window after co-selection for each locus. Values represents the fraction of reads with either one, two or three edits compared to the total fraction of reads that were edited. **D)** Editing outcome type for all sites with a total editing rate greater than one percent after co-selection (n=30 cytosines across all targeted sites). The C to G/T distribution represents the sum of editing that resulted in a C to G or C to T mutation. Position-wise editing rates and outcome are shown in Supplementary Figures 5 and 6. **E)** Agreement between the predicted nucleotide total editing rank in the model used to predict mutagenesis outcomes in the large-scale experiment and the deep sequencing data (n=28 sites, 10 gRNAs: gRNA specific predicted and observed rankings are presented in Supplementary Figure 5 and 6). The gRNAs targeting *ADE1* and *SES1* were respectively excluded from the analysis because there is only one editable site in the activity window and total editing rate was too low. **F)** Edited read coverage of the mutation outcome prediction model and the 99th percentile of edited allele combinations (n=4 genotypes in both cases) for the gRNAs with editing activity included in the large-scale experiment.

In our editing model, we first predict that single mutants would be the main mutagenesis outcome of the base editing process. We found this to be true for 9 gRNAs out of 10 with more than one cytosine in the Target-AID activity window (Figure 1C). Second, our model considers C to A editing to be rare and thus disregards them in favor of the more common C to G and C to T mutations. We observe this bias in the deep sequencing data (Figure 1D), with the median occupancy of both C to G and C to T genotypes in edited alleles being much greater than C to A occupancy (C to T vs C to A: *W*=0, p=1.73×10^-6^, C to G vs C to A: *W*=41, p=8.19×10^-5^, two-sided wilcoxon signed rank test). Including these mutations as in our model leads to a median coverage of 93% of mutagenesis outcomes. Our sequencing data also showed a greater prevalence of C to T mutations compared to C to G (*W*=112, p=0.01), but if absolute editing rate is taken into account this difference disappears (Supplementary Figure 4F). Finally, in cases where multiple editable nucleotides are present in the activity window of the base editor, our model uses the quantitative data of the original Target-AID manuscript to predict qualitatively which position should be edited at the highest frequency. We found that this prediction method of editing rank in the activity window matched with the experimental data in most cases (Figure 1E) which is unlikely to occur by chance (p≈ 0.0004 based on 1×10^6^ random rank permutations). Globally, we found that the edited allele pool was mostly composed of the genotypes predicted by our model: for the 8 gRNAs with editing activity that came from the library, the median fraction of edited reads covered by our model was 69% (Figure 1F). In 7 out of 8 cases, the fractions of edited reads covered by the model was better than the 99th percentile of randomized outcome combinations and in 6 out of 8 cases and also superior to the 99.9th percentile. Overall, these results support that a large fraction of the gRNAs included in our library can edit their genomic targets in an efficient and predictable manner.

### High throughput screening using the gRNA library

The gRNA library was cloned into a high-throughput co-selection base editing vector^23^. We performed pooled mutagenesis followed by bulk competition (Supplementary Figure 7) to identify mutations with significant fitness effects (Figure 2). As the relative abundance of each gRNA in the extracted plasmid pool depends on the abundance of the subpopulation of cells bearing these gRNAs, any fitness effect caused by the mutation they induce will influence their relative abundance. Variation in plasmid abundance was measured using targeted next-generation sequencing of the variable gRNA locus on the base editing vector in a manner similar to GeCKO approaches^6, 26^.

**Figure 2.**
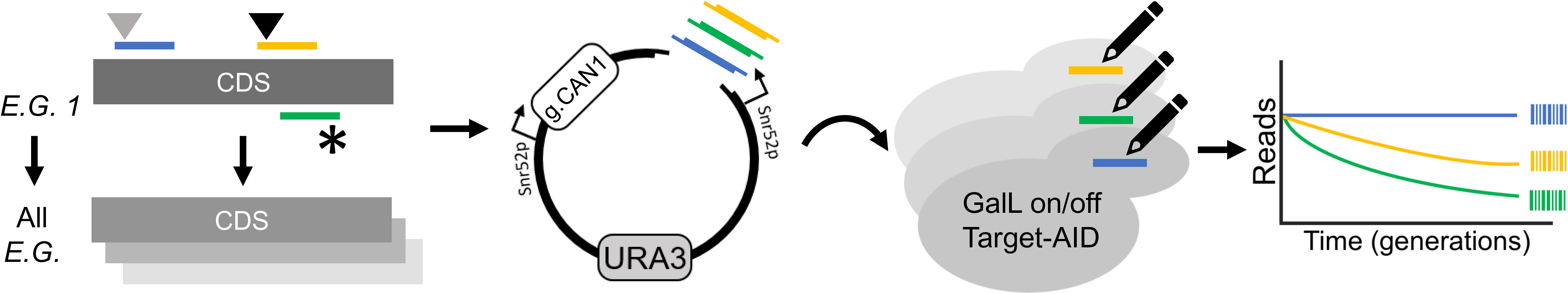
A gRNA library for systematic perturbation of essential genes using the Target-AID base editor. Essential genes (ex.: *E.G.1*) were scanned for sites appropriate for Target-AID mutagenesis. Mutational outcomes include silent (grey triangle), missense (black triangle) mutations, as well as stop codons (*). DNA fragments corresponding to the gRNA sequences were synthesized as an oligonucleotide pool and cloned into a co-selection base editing vector. Using gRNAs as molecular barcodes, the abundance of cell subpopulations bearing mutations is then measured after mutagenesis and bulk competition. Mutations with fitness effects are inferred from reductions in the relative gRNA abundances.

After applying a stringent filtering threshold based on gRNA read count at the mutagenesis step (see methods), we identified a total of ∼17,000 gRNAs for which we could evaluate fitness effects. Replicate data for gRNAs passing the minimal read count selection criteria showed high correlation across experimental time points (Supplementary Figure 8) and cluster by experimental step (Supplementary Figure 9), showing that the approach is reproducible. Using the distribution of abundance variation of gRNAs between the start of the screen and the end of mock glucose induction as null distribution, we identified 1,118 gRNAs across 605 loci with significant negative effects (GNE) on cell survival or proliferation at a 5% False Discovery Rate (Figure 3A, Supplementary Figure 9 B and C). GNEs are distributed evenly across the yeast genome (Figure 3B), suggesting no inherent bias against specific regions. An example of gRNA abundance variation through time for all gRNAs (both GNEs and NSGs) targeting *GLN4* is shown in Figure 3C.

**Figure 3.**
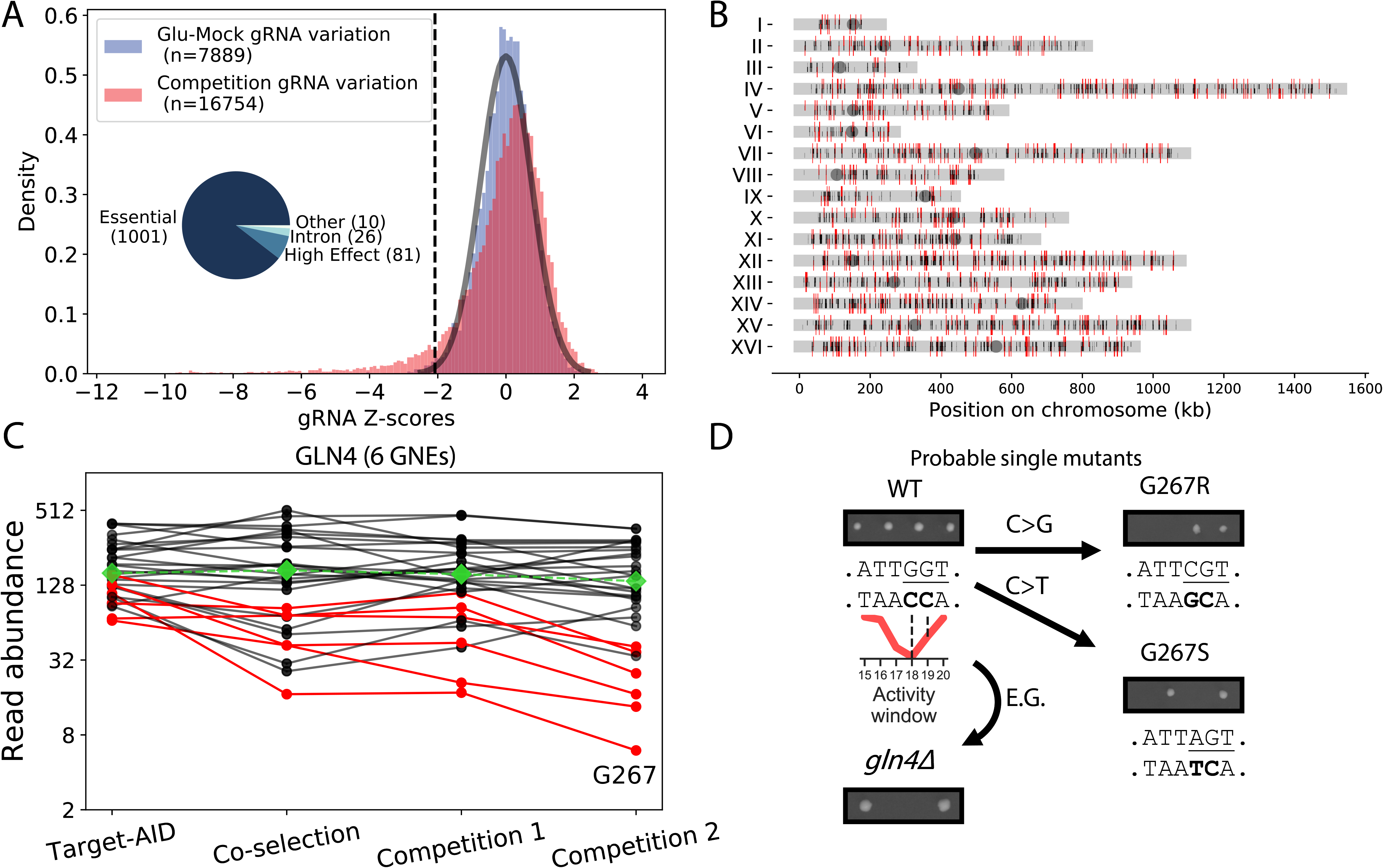
High-throughput forward mutagenesis by Target-AID base editing identifies sensitive sites across the yeast genome. **A)** Cumulative distribution of z-scores of the log2 fold-change in gRNA abundance between mutagenesis and the end of the bulk competition experiment. Scores were calculated using the distribution of abundance variation between the start of the experiment and the end of mock editor induction, the fitted normal distribution is shown as a black line. The z-score threshold was set at ∼5% FDR and is represented by a dotted black line. The distribution of target types in the 1,118 gRNAs with Negative Effects (GNE) is shown in the inset. **B)** Positions of base editing target sites in the yeast genome. Telomeric regions are depleted in target sites because very few essential genes are located there. GNEs are shown in red, and other gRNAs are in black. The orientation of the line matches the targeted strand relative to the annotated coding sequence. **C)** Decline in gRNA abundance (on a log scale) between timepoints after mutagenesis for gRNAs targeting GLN4, a tRNA synthetase. Median gRNA abundance across the entire library through time is shown in green. The red lines represent the gRNAs categorized as having a significant effect (GNE) for this gene, while non-significant gRNAs (NSG) are shown in black. The gRNA with the most extreme z-score targets residue G267. **D)** Mutagenesis of GLN4-G267 confirms its essential role for protein function. Tetrad dissection of a heterozygous deletion mutant bearing an empty vector results in only two viable spores, while the wild-type copy in the same vector restores growth. Dissection of the two heterozygous mutants bearing a plasmid with the most probable single mutant based on the known activity window of Target-AID shows both mutations are lethal.

Because our screen specifically targeted essential genes, many gRNAs cause mutations in highly conserved regions with high functional importance. To illustrate this, we focus on the highest scoring GNE targeting *GLN4*, a tRNA synthetase. The gRNA 33725 mutates a glycine at position 267 into either arginine or serine, and showed a dramatic drop in abundance in the large-scale experiment. To validate the deleteriousness of the predicted mutations, we transformed a centromeric plasmid bearing a wild-type or mutated copy of the gene under the control of its native promoter^27^ in a heterozygous deletion background^28^ (Supplementary figure 10A). Glycine 267 is part of the “HIGH” motif, characteristic of class I tRNA synthetases, and is involved in ATP binding and catalysis and is highly conserved through evolution^29^. As expected, the region around the “HIGH” motif shows both a low evolutionary rate based on inter-species comparisons and a much lower variant density in yeast populations compared to other domains of Gln4 (Supplementary figure 10B), showing conservation both on a short and long timescales. Surprisingly, mutagenesis experiments in the bacterial homolog MetRS concluded that mutating this residue from glycine to alanine did not alter significantly catalysis while mutating it to proline had a strong disruptive effect^30^. We found that mutating Gly 267 either to Arg or Ser was enough to cause protein loss of function (Figure 3D).

The five other sensitive sites identified in GLN4 by our screen were also clustered in regions with slow evolutionary rates. We found that one other GNE targeting residue D291 induced a highly deleterious mutation coupled with a neutral mutation as outcomes (D291E vs D291D, Supplementary Figure 11). We did not observe any discernible growth defect for the other GNE outcomes and as well as for the outcomes of 4 NSG targeting nearby amino acids. The other GNEs tested had markedly more positive scores than the one targeting G267, which would be consequent with a higher false positive rate close to the significance threshold. However, the case of the D291E/D291D pair, where a strong fitness effect is partially obscured by a neutral mutation produced by the other mutagenesis outcomes supports that sites of interest can be detected even close to the significance threshold. As we only tested two outcomes per gRNA, it is also possible that some of the abundance drops we measured were the result of mutations outside of our model, which are sometimes predicted to be more deleterious than the most likely mutations.

### Comparison of GNE induced mutations with variant effect predictions

If GNEs indeed induce specific deleterious mutations, these mutations should be predicted to be more deleterious than those of Non-Significant gRNAs (NSG). We tested this using two recently published resources for variant effect prediction: Envision^2^ and Mutfunc^31^. Envision is based on a machine learning approach that leverages large-scale saturated mutagenesis data of multiple proteins to perform quantitative predictions of missense mutation effects on protein function. The lower the Envision score, the higher the effect on protein function. Mutfunc aggregates multiple types of information such as residue conservation through the use of SIFT^32^ as well as structural constraints to provide a binary prediction of variant effect based on multiple quantitative and qualitative values. Mutations with a low SIFT score have a lower chance of being tolerated, while those with a positive ΔΔG are predicted to destabilize protein structure or interactions. Both Envision and the Mutfunc aggregated SIFT data cover the majority of the most probable mutations generated by the gRNA library (Supplementary Figure 12A). The structural modeling information had much lower coverage, covering at best around 12% of the most probable mutations (Supplementary Figure 12B). As expected, mutations generated by GNEs showed significantly lower SIFT scores and showed enrichment for strong effects predicted by SIFT and Envision (Figure 4). Indeed, all four most probable substitutions created by GNEs are about twice more likely to be predicted to have a large deleterious effect by Envision or a very low chance of being tolerated as predicted by SIFT compared to NSG gRNAs. Envision scores across the proteome show a high level of homogeneity, with most mutations having a score between 0.94 and 0.96 (Supplementary Figure 12C). According to the original Envision manuscript, this should be predictive of a small decrease in protein function. As such, the shifts in score distributions between GNEs and NSGs are more subtle but still support that GNE induced mutations are generally more likely to be deleterious as well (Supplementary Figure 13A).

**Figure 4:**
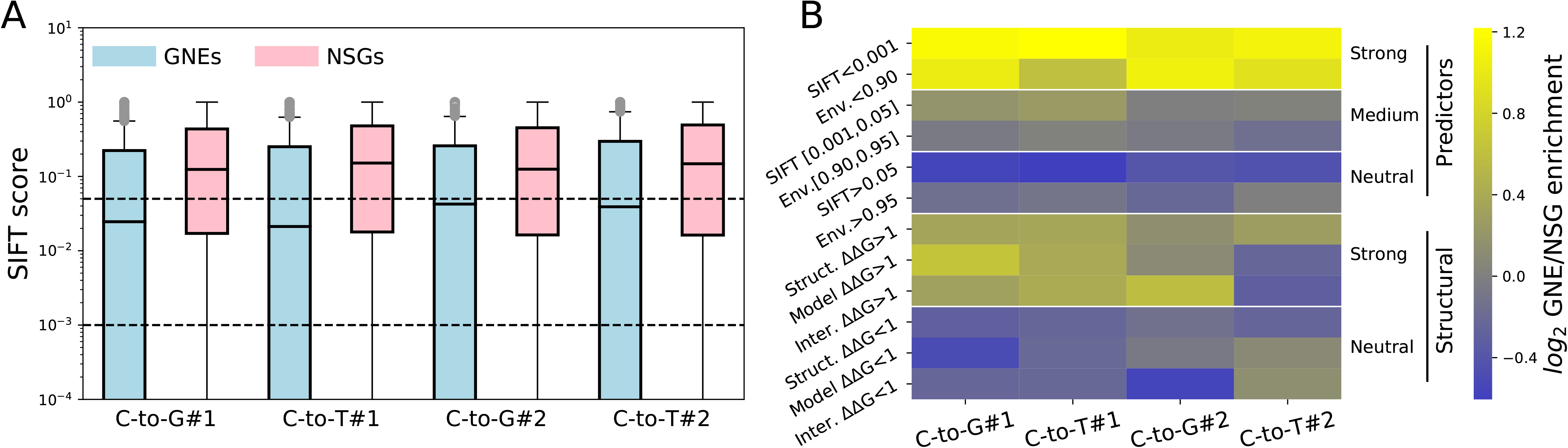
GNE induced mutations are enriched in predicted deleterious effects. **A)** SIFT score distributions for the most likely induced mutations of both GNEs (blue) and NSGs (red). The thresholds for the categories used in the enrichment calculations in **B)** are shown as black dotted lines. SIFT scores represent the probability of a specific mutation being tolerated based on evolutionary information: the first threshold of 0.05 was set by the authors in the original manuscript^32^ but might be permissive considering the number of mutations tested in our experiment (n= 895, 12394, 704, 8520, 643, 7396, 508, 5682). All GNE vs NSG score comparisons are significant (Welch’s t-test p-values: 1.19×10^-24^, 3.01×10^-24^, 9.00×10^-12^, 1.55×10^-12^). The box cutoff is due to the large fraction of mutations for which the SIFT score is 0. B) Enrichment folds of GNEs over NSGs for different variant effect prediction measurements. Envision score (Env.), SIFT score (SIFT), protein folding stability based on solved protein structures (Struct. ΔΔG), protein folding based on homology models (Model ΔΔG) and protein-protein interaction interface stability based on structure data (Inter. ΔΔ G). The raw values used to calculate ratios are shown in Supplementary table 1. The predictions based on conservation and experimental data are grouped under ‘Predictors’ and those based on the computational analysis of protein structures and complexes under ‘Structural’.

Mutations with destabilizing effects as predicted by structural data also appeared to be enriched in GNEs predicted mutations but low residue coverage limits the strength of this association. This is supported by the raw ΔΔG value distributions, which show a significant tendency for GNE mutations to be more destabilizing (Welch’s t-test p-values for GNE vs NSG ΔΔG: C-to-G #1 0.0001, C-to-T #1 0.0064, C-to-G #2 0.148, C-to-T #2 0.007, Supplementary Figure S13B-D). However, the shift in distribution only achieved significance for certain mutation predictions based on solved structures and homology models. While low residue coverage limits our statistical power, this weak apparent enrichment for mutations affecting protein stability may having a limited effects on fitness. As expected from known experimental data on mutagenesis outcomes^15^, signal was usually stronger for the most probable C to G mutation.

### Sensitive sites provide new biological insights

Since Target-AID can only generate a limited range of amino acid substitutions from a specific coding sequence, we investigated whether any of these mutational patterns were enriched in GNEs (Figure 5A, source data in Supplementary tables 2, 3, and 4). We found deviations from random expectations in both C-to-G and C-to-T mutation ratios that drove the enrichment of several mutation combination. Three out of four of the mutation pair patterns involving glycine were enriched in GNEs. For example, the Glycine to Arginine or Serine substitutions (as exemplified by guide 33725 targeting *GLN4*) is the second most enriched pattern, being almost four-fold overrepresented in GNE outcomes. This pattern is consistent with the fact that Arginine has properties highly dissimilar to those of Glycine^34^, making these substitutions highly deleterious. Furthermore, as Glycine residues are often important components of cofactor binding motifs (eg.: Phosphates)^35^ this observation might reflect a tendency for GNEs to alter these sites.

**Figure 5.**
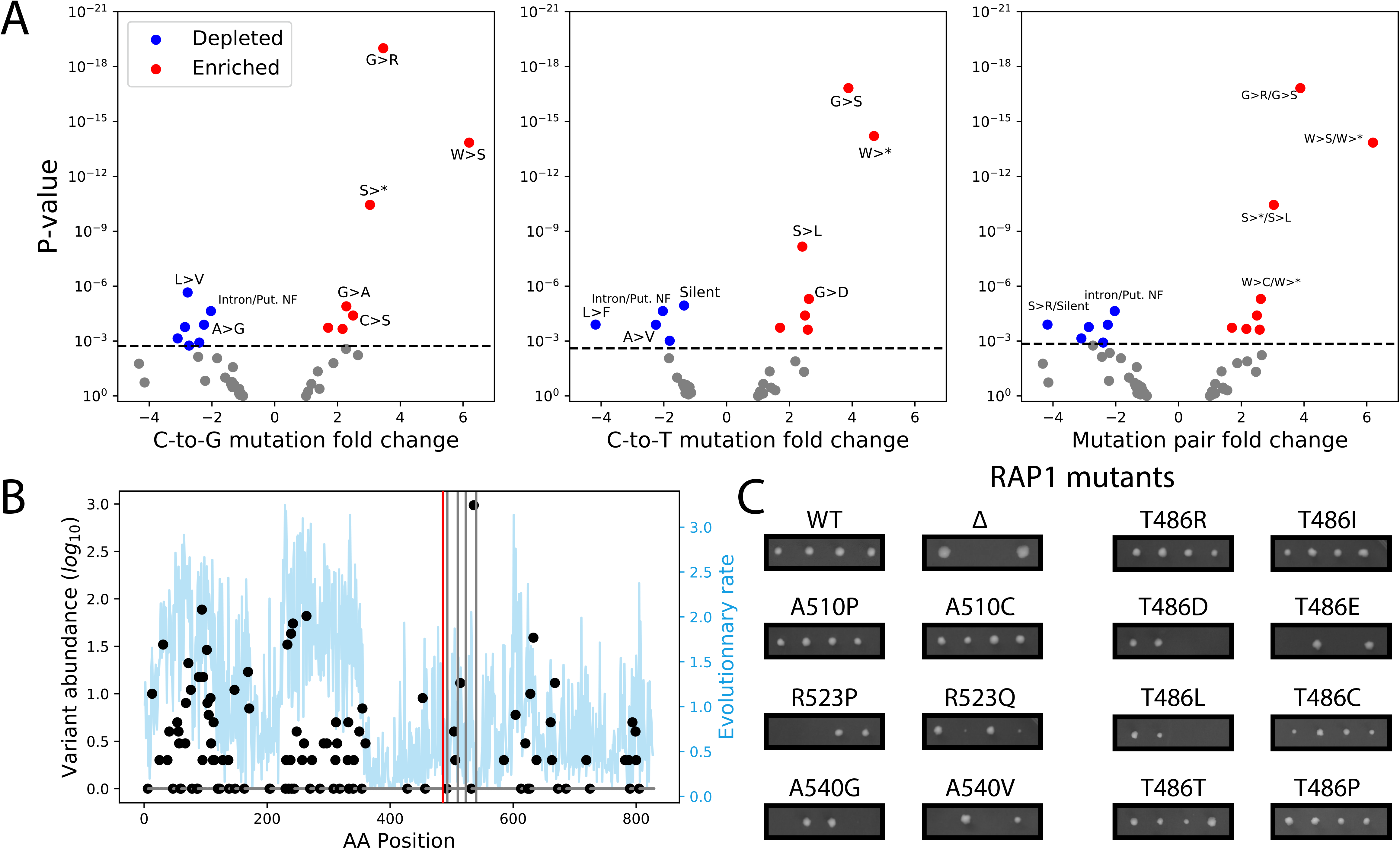
GNE mutations are enriched for specific amino acid substitution patterns and identify critical sites for protein function. **A)** Fold depletion and enrichment volcano plots for the most probable mutations induced by GNEs in the screen. Enrichment and depletion values were calculated by comparing the relative abundance of each mutation among GNEs and NSGs using Fisher’s exact tests. Mutation patterns significantly depleted are shown in blue, while those that are enriched are in red. The significance threshold was set using the Holm-Bonferroni method at 5% FDR and is shown as a dotted grey line. **B)** Protein variant frequency among 1000 yeast isolates (black dots) and residue evolutionary rate across species (blue line) for *RAP1*. The target site for the GNEs targeting T486 is highlighted by a red line while the other detected GNEs target sites are shown by a grey line. **C)** Tetrad dissections confirm most *RAP1* GNE induced mutations indeed have strong fitness effects, as well as other substitutions targeting these sites.

As expected, there is a strong enrichment within GNEs for patterns that result in mutation to stop codons: both C-to-G patterns (Y to stop: 3 fold enrichment, p=3.62×10^-11^, S to stop: 2.2 fold enrichment, p=0.0002) but only one C-to-T pattern was overrepresented significantly (W to stop, 4.6 fold enrichment, p=6.23×10^-15^). Substitutions to stop codon in one outcome also drove enrichment in the other: for example, the link between Serine to Stop (C-to-G) appears to be the cause of the Serine to Leucine (C-to-T) overrepresentation. Both mutation pairs involving mutating a Tryptophan to a stop via a C-to-T mutation are enriched: this is not surprising, as the alternative mutations Tryptophan to Serine or Cysteine are also highly disruptive^34^. Changes between similar amino acids, which are expected to be tolerable, were also generally depleted in GNE (ex.: the Alanine to Glycine/Valine pair). Mutations in intronic sequences and putative non-functional peptides were also underrepresented, as were most patterns leading to silent mutations (Figure 5A). These results show the power of this approach to discriminate important functional sites from more mutation tolerant ones across the genome.

Interestingly, genes for which more than one GNE were detected were enriched for molecular function terms linked to cofactor binding (Supplementary Table 5). This suggests that the GNEs might indeed have a tendency to affect protein function through mechanisms other than protein or interaction interface destabilization. These protein properties depend on many residues, making them more robust to single amino acid substitutions, whereas cofactor binding may depend specifically on a handful of residues, making these sites critical for function. Using the Uniprot database^37^, we also examined whether gRNAs that target annotated binding sites or highly conserved motifs are more likely to affect fitness compared to other gRNAs targeting the same set of genes. We found a 3.5 fold enrichment for GNEs directly affecting these sites (49/188, ratio^GNE On^=0.261, 447/5969 ratio^GNE Off^=0.0749, two-sided Fisher’s exact test p=3.54×10^-14^) or residues in a two amino acid window around them (23/138, ratio^GNE near^=0.167, 447/5969, ratio^GNE Off^=0.0749, two-sided Fisher’s exact test p=0.00048).

The precise targeting of our method also allows us to investigate amino acid residues with known functional annotations such as post-translational modifications. We found no significant enrichment for gRNAs mutating directly annotated PTMs (ratio^GNE PTM^ = 19/1118, ratio^NSG PTM^ 243/15536, Fisher’s exact test p=0.71). Most of these sites were phosphorylation sites (7), metal coordinating residues (5) and ubiquitination sites (4). This is consistent with the hypothesis that many PTM sites may have little functional importance^36^ and thus mutations affecting them should not be significantly enriched for strong fitness effects compared to other possible mutations. The same was also observed for gRNAs mutating residues near known PTMs that could disturb recognition sites (ratio^GNE nearPTM^ = 130/1118, ratio^NSG nearPTM^ = 1698/15536, Fisher’s exact test p=0.43). As we did not specifically target PTMs, our sample size is small and it should be noted that statistical power regarding these observations is limited.

However, GNEs that do target annotated PTM sites might provide additional evidence supporting the importance of these sites in particular. For example, the best scoring GNE in the well-studied transcriptional regulator *RAP1* is predicted to mutate residue T486. This threonine has been reported as phosphorylated in two previous studies^38, 39^, but the functional importance of this phosphorylation has not been explored yet. Residue T486 is located in a disordered region in the DNA binding domains^40^, which part of the only *RAP1* fragment essential for cell growth^41, 42^. Because the available wild-type *RAP1* plasmid (see methods) does not complement gene deletion growth phenotype, we used a different strategy for validation that relied on CRISPR-mediated knock-in (see methods and Supplementary Figure 14). We tested the effect of several predicted GNE induced mutations in RAP1 targeting positions T486, A510, R523 and A540 (Figure 5B-C). We found that the predicted mutations at two of these positions, R523 and A540, were highly deleterious. While we could not validate that the two most likely mutations predicted to be caused by the GNE targeting T486 had a detectable fitness effect in these conditions, we found that phosphomimetic mutations at this position were lethal but most other amino acids were well tolerated. While we could validate that this gRNA indeed targeted a sensitive site, the outcomes predicted by our model did not have any detectable fitness effects. This showcases a limitation of our approach: the uncertainty in outcome prediction can complicate validation studies. As we only tested progeny survival on rich media and at a permissive temperature and the screen was performed in synthetic media at 30°C, these mutants might still affect cell phenotype but in an environment-dependent manner.

### gRNA properties influence mutagenesis efficiency

There are still very few high-throughput experimental datasets available that allow the investigation of which gRNA properties affect editing efficiency in the context of base editing. We therefore sought to examine what gRNA and target sequence features could influence mutagenesis efficiency. To do so, we focused on the subset of gRNAs with the potential to generate stop codons (stop codon generating gRNAs, SGGs) in essential genes (Figure 6A). As gRNAs in our library were designed to target the first 75% of the coding sequences, successful stop codon generation in this subset of genes should often lead to a lethal loss of function^13, 22^.

**Figure 6.**
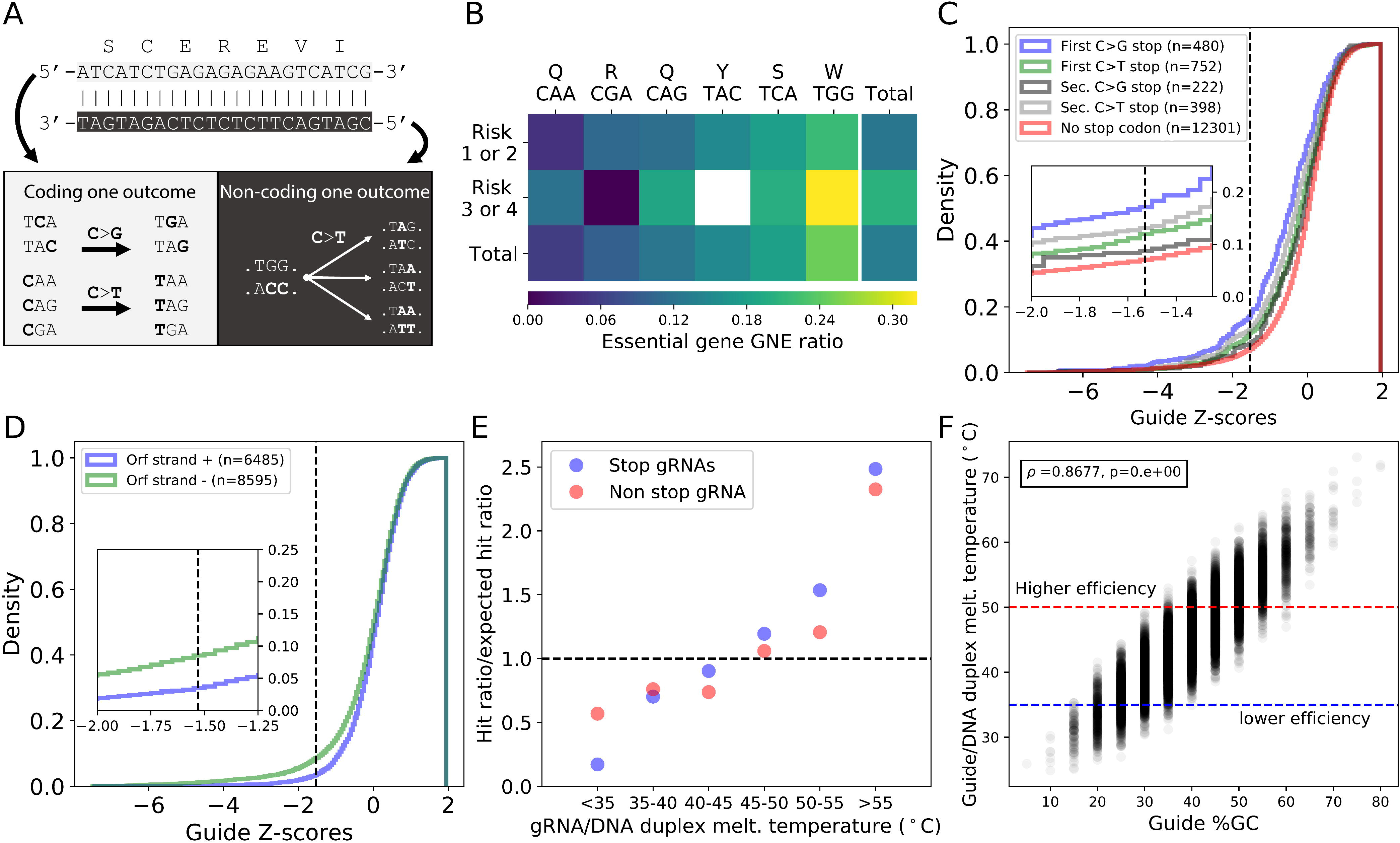
gRNA and target properties affect mutagenesis efficiency. **A)** Since Target-AID can generate both C to G and C to T mutations, many codons can be targeted to create premature stop codons. The TGG (W) codon is the only one targeted on the non-coding strand as ACC. **B)** GNE ratio for SGGs targeting different codons in essential genes, split by co-editing risk categories, were 1 and 2 represent low or very low co-editing risk while 3 or 4 represent moderate to high co-editing risk. **C)** Cumulative z-score density of SGGs grouped by the mutational outcome generating the stop codon. A higher rate of GNE is observed for gRNAs for which a C-to-G mutation at the highest editing activity position generates a stop codon mutation. The significance threshold is shown as a black dotted line. **D)** Cumulative z-score density of gRNAs that do not generate stop codons targeting either the coding or non-coding strand. **E)** SGG and non-SGG GNE enrichment compared to the expected GNE ratio for different melting temperature ranges. **F)** gRNA/DNA duplex melting temperature as a function of gRNA GC content for all gRNAs for which fitness effects were measured. The higher and lower efficiency thresholds are based on the enrichments shown in panel E.

We found important variation in the ratio of GNE for the different types of SGGs (Figure 6B), with gRNAs targeting TGG (Trp) codons having the highest activity. This is in opposition to the general trend, as in general C to G mutation leading to stop codon formation had higher GNE ratios than the three other C-to-T alternatives. Overall, we observed significant GNE enrichment in SGGs which depend on the first C to G mutation to induce stop codon formation (Figure 6C). Multiple factors can explain the higher performance of TGG targeting gRNAs. First, as most of these sites have high co-editing risk scores because of the two consecutive cytosines, they might have increased editing rates due to processive co-editing events, increasing the chance of fitness effect detection. This phenomenon might also occur in non-SGG gRNAs (Supplementary Figure 15A). Second, we found a significant enrichment in GNEs for gRNAs targeting the non-coding strand, even after excluding SGGs (Figure 6D). This effect might be explained by the higher repair efficiency in the transcribed strand in yeast^43^. Furthermore, as the non-coding strand is the one which is transcribed, a deamination event there might lead to consequences at the protein level more rapidly when the mutated coding sequence is transcribed. In contrast, the targeted chromosomal strand appears to be much less important (Supplementary Figure 15B). The variation in GNE ratio observed between the different SGG target codons might also reflect *in vivo* DNA repair preferences that depend on sequence context, where different outcomes might be favored depending on the target sequence. For example, the CA di-nucleotide might favor C to G mutations, which would explain the low GNE ratio of CAA (Gln) targeting SGGs and the higher than average GNE ratio of TCA (Ser) targeting SGGs.

Another parameter with a high impact on GNE enrichment in gRNA sets is the predicted melting temperature of the RNA-DNA duplex formed by the gRNA sequence and its target DNA sequence (Supplementary Figure 15C-D). Both SGG and non-SGG gRNAs with low values have a lower chance of being detected as having effects, while gRNAs with higher values are enriched for GNEs (Figure 6E). This enrichment cannot be attributed to technical biases in library preparation or high-throughput sequencing that would tend to lower their abundance as melting temperature shows practically no correlation with read count at any time point (Supplementary Figure 16). Furthermore, this effect is not caused by target position bias within target genes or a strong correlation between GC content and the targeted position (Supplementary Figure 17). Even if binding energy is strongly correlated with GC content, there is still significant variation within gRNA sets with the same %GC (Figure 6F).

## Discussion

Using targeted deep sequencing and high throughput screening, we investigated whether the Target-AID base editor is amenable for genome-scale targeted mutagenesis studies. We show that a prediction model based on known Target-AID properties can be used to predict the major mutational outcome of editing, even if multiple editable nucleotides are present in the activity window. Using yeast essential genes as a test case, we then applied this approach on a larger scale and identified hundreds of gRNAs targeting sensitive residues that have significant effects on cellular fitness when mutated. We could then verify orthogonally the effects of mutational outcomes of GNE using classical genetics approaches and show that they tend to overlap with variants predicted to be deleterious. By focusing on a few highly relevant variant sets, we highlighted the power and potential of our approach to generate new biological insights. We then used this data to investigate which factors influence base editing efficiency and found multiple gRNAs and target properties that affect mutagenesis and that could be optimized for future experiments in specific genomic spaces.

In previously published methods such as TAM and CRISPR-X^18, 19^, the semi-random nature of the editing forces the use of mutant allele frequencies as a readout for mutational fitness effects, potentially limiting the scale of the experiments because only one genomic region can be targeted at a time. To complement these approaches, we use more predictable base editing to increase dramatically the number of target loci, albeit at the cost of a lower mutational density. Our results demonstrate the feasibility of base editing screening at a large scale with applications beyond stop codon generation, and future developments will further enhance it. For instance, the use of a base editor with multiple possible mutagenesis outcomes complexifies the prediction of editing outcomes, which can, in turn, make GNE follow-up challenging. Using a base editor that channels mutational outcomes such as cytidine deaminase-uracil glycosylase inhibitor (UGI) fusion can address this problem^15^ but decreases the number of mutations explored during the experiment. However, recently published data on cytidine deaminase-UGI fusion has shown they could lead to off-target editing in vivo at a much higher rate compared to adenine base editors or the Cas9 nuclease^44, 45^. Although there is currently no high throughput data on the off-target activity of Target-AID, data generated in yeast in the original publication suggests far lower rates than those recently reported in mammalian cells^15^. Recently, Sadhu, Bloom et al examined the effects of premature stop codons (PTC) in essential genes using a high throughput variant construction method that relied on homology directed repair using a mutated repair template^13^. They observed that a significant fraction of PTCs can be tolerated, but only within the last 30 codons of a protein. Outside this window, they found no link between PTC tolerance and position within the coding sequence, something which we also did not observe both for SGGs and non-SGG gRNAs (Supplementary Figure 17A-B).

We provide key empirical data on gRNA dependant parameters that can be used to optimize base editing efficiency. Based on our results, selecting gRNAs with high binding energy to their genomic targets and favoring those which target the non-coding strand can increase the chance of high editing activity. Importantly, our observations differ from what has been reported for Cas9-based genome editing. High gRNA RNA/DNA duplex binding has instead been associated with lower mutagenesis efficiency^46^. Our data thus confirms the observation that parameters associated with Cas9 editing cannot readily be transferred to base editors^47^. Furthermore, the temperature at which experiments are performed might affect efficiency for certain gRNAs with low gRNA-DNA duplex binding energy and should be considered when designing base editing experiments in different organisms^15^. However, it remains to be confirmed whether the enrichment for certain gRNA properties we observed are specific to Target-AID or will also be transferable to other base editors as this may depend on the enzymatic properties of these proteins. Acquiring large paired gRNA and mutagenesis outcome datasets similar to those available for Cas9 genome editing^24^ will allow for more refined models for rational base editing activity prediction.

The field of base editing is rapidly evolving, with new tools being developed constantly. One of the most recent additions to this fast-growing toolkit are engineered Cas9 enzymes with broadened PAM specificities^48^, which have already been shown to be compatible with base editors. More flexible PAM requirements are especially useful for base editing applications, as they increase the number of sites to be edited and also the number of potential gRNAs per site, increasing the chances of choosing optimal properties and thus greater efficiency^25^. Our method allows an experimental scale which bridges saturation mutagenesis methods and genome-wide knock-out studies, alleviating the current trade-off between mutational diversity and the number of targets genes to generate new biological insights.

## Methods

### Generation of a gRNA library for Target-AID mutagenesis of essential genes in yeast

The Target-AID base editor has an activity window between base 15 to 20 in the gRNA sequence starting from the PAM, and the efficiency at these different positions was characterized in Nishida *et al.* 2016. This allowed us to predict the mutational outcomes for a specific gRNA provided the number of editable bases in the window is not too high. To select gRNAs, we parsed a database of gRNA targets for the *S. cerevisiae* reference genome sequences (strain S288c)^49^ and applied several selection criteria. Since the screen was to be performed in the BY4741 strain, all gRNAs (unique seed sequence, no NAG site) within the database were aligned to the reference genome of that strain using Bowtie^50^. Only gRNAs with a single perfect alignment were kept for subsequent steps. To select gRNAs amenable to Target-AID base editing, we selected gRNAs with cytosines within the highest activity window of the editor (positions −17 to −19 starting from the PAM). To limit the total number of possible mutational outcomes, gRNAs with three cytosines within the window were removed as well as those with two cytosines at the highest activity positions. Next, we filtered out any gRNA containing a BsaI restriction site to prevent errors during the library cloning step.

The list of essential genes (n=1156)^3, 4^ was used to discriminate between gRNAs targeting essential or non-essential genes (retrieved from http://www-sequence.stanford.edu/group/yeast_deletion_project/Essential_ORFs.txt). Among non-essential genes, data from Qian et al. 2012^51^ was used to create categories of fitness effects. If the fitness score (averaged across media and replicates) of a gene was below 0.75, it was categorized as “high effect” on fitness. We excluded auxotrophic marker genes as well as *CAN1*, *LYP1*, and *FCY1* because those could be used as co-selection markers^23^. Gene deletions with an averaged fitness score between 0.999 and 1.001 were categorized as having “no detectable effect” on fitness. We selected gRNAs targeting essential and high effect genes, as well as gRNAs targeting a set of 38 randomly chosen no effect genes. To further limit the space of gRNAs examined, only gRNAs mapping from the 0.5^th^ percent to the 75^th^ percent of coding sequences were chosen. We also added gRNAs targeting all known yeast introns (Ares lab Database 4.3)^52^ and putative non-functional peptides^53^ selected with the same strategy except for the constraints on gRNA position within the sequence of interest. This resulted in a set of 39,989 gRNAs: library properties are summarized in Supplementary Figure 1. To assign a co-editing risk score to each gRNA, we defined four categories using the extended activity window sequence composition shown in Table 1.

**Table 1:**
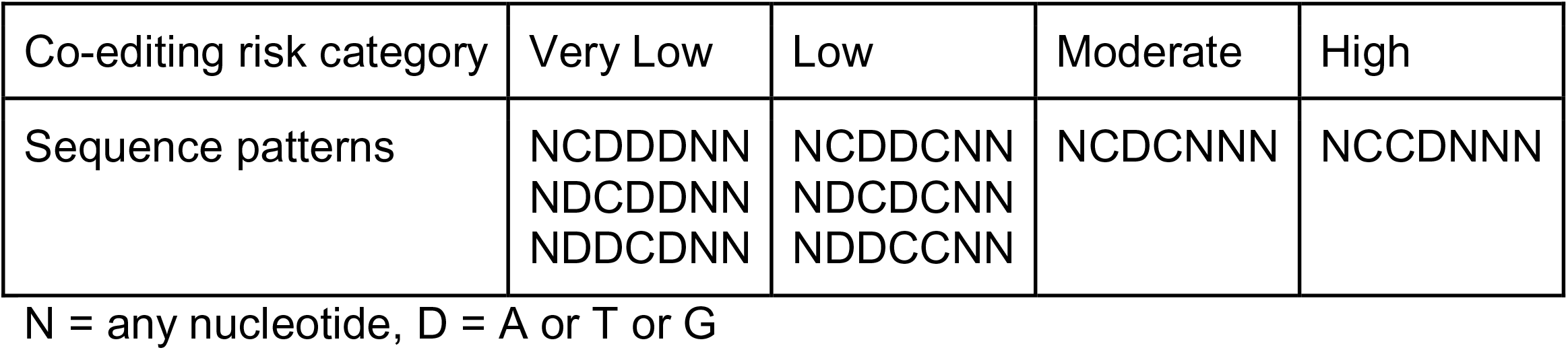
Sequence patterns of co-editing risk categories.

### Library construction

The plasmids, oligonucleotides, and media used in this study are listed in as Supplementary tables 6, 7 and 8 respectively. The oligo pool was synthesized by Arbor Biosciences (Michigan, USA) and was cloned into the pDYSCKO vector using Golden Gate Assembly (New England Biolabs, Massachusetts, USA) with the following reaction parameters:

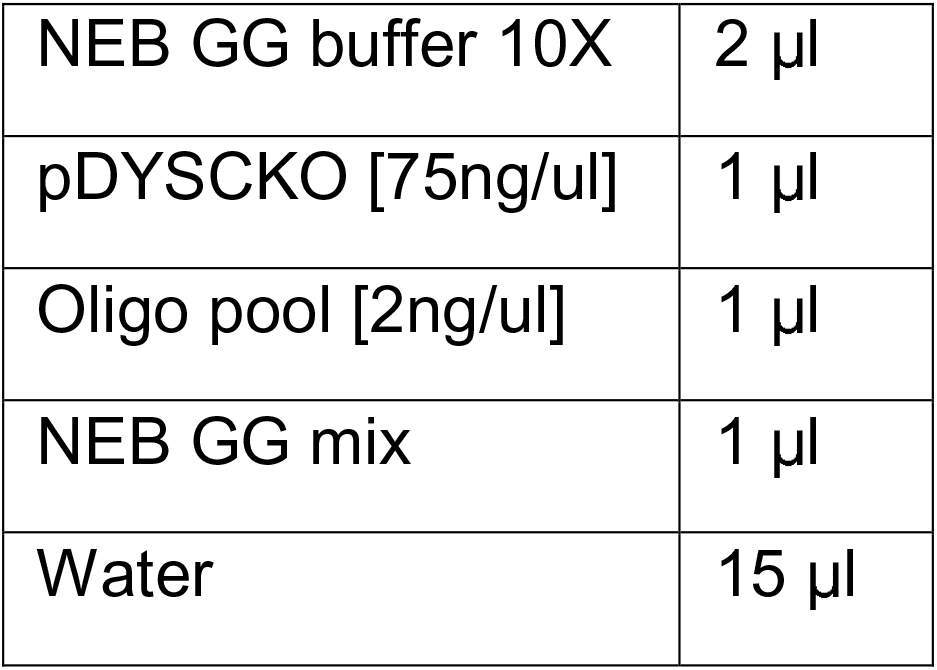

The ligation mix was transformed in *E. coli* strain MC1061 ([*araD139*]*_B/r_* Δ*(araA-leu)7697* Δ*lacX74 galK16 galE15(GalS)* λ*-e14-mcrA0 relA1 rpsL150(strR) spoT1 mcrB1 hsdR2*)^54^ using a standard chemical transformation protocol and plated on ampicillin selective media to select for transformants. Serial dilution of cells after outgrowth were plated and then used to calculate the total number of clones produced by the cloning reaction. Quality control of the assembly was performed by Sanger sequencing ∼10 clones per assembly reaction. Cells were scraped from plates by adding ∼5 ml of sterile water, incubating a few minutes at room temperature, and then using a glass rake to resuspend colonies. Resuspended plates were then pooled together in a single flask per reaction, which was then used to make glycerol stocks of the library and cell pellets for plasmid extraction. The Qiagen Midi-Prep kit (Qiagen, Germany) was used to extract plasmid DNA from cell pellets by following the manufacturer’s instructions. The DNA concentration of each eluate was then measured using a NanoDrop (Thermofisher, Massachusetts, USA), and a normalized master library for yeast transformation was assembled by combining equal quantities of each assembly pool.

### Base editing time course and library preparation for deep sequencing

Cells were co-transformed with pKN1252 and the pDYSCKO plasmid bearing the gRNA of interest using the protocol described below for the large-scale experiment. Transformant plates were scraped by adding ∼5 ml of sterile water, incubating a few minutes at room temperature, and then using a glass rake to resuspend colonies. The resuspended cells (one pool per guide) were used to inoculate two replicate cultures per guide. Cells went through the same induction protocol as for the large-scale experiment, but scaled down to a 24 deepwell plate (see Supplementary Figures 3 and 7). The volumes used were: 3 ml for the initial SC-UL+glucose culture, 4 ml for the SC-UL+glycerol step, 3 ml for the SC-UL+galactose step, and 3 ml for the liquid canavanine co-selection step. At the end of the galactose induction step, 100 μ 1/2000 dilution of each well was plated on SC-ULR+canavanine solid media to obtain editing survivor colonies. At the glycerol to galactose media switch, a ∼1 OD pellet was sampled by spinning cells at 13 200 RPM and removing the media. Cell pellets were then stored at −80°C for subsequent DNA extraction. The same method was used to sample ∼1 OD at T=6 hours in galactose, ∼2 OD at T=12 hours in galactose, and ∼3 OD at the end of canavanine co-selection. Plates with selected colonies (edited at the CAN1 locus) were soaked in water and scraped, and 1.4 ml of the resulting cell suspension was sampled and stored.

Genomic DNA was extracted from cell pellets using a standard phenol-chloroform method from each sample^55^ and quantified by NanoDrop (Thermo fisher, Massachusetts, USA). For each sample, we aimed to sequence both the target edit site and the *CAN1* co-selection edit site. To multiplex the 240 samples in the same sequencing library, we used the row-column-plate-indexed PCR (RCP-PCR) approach^56^. Briefly, each target locus was amplified from genomic DNA and universal adapter sequences were added to each end of the amplicon. A 1/2500 dilution of the resulting product was then used as template with a set of 10 (rows) by 12 (column) primers used to index each sample in a second PCR reaction. All samples for the same locus were then pooled together and normalized according to electrophoresis gel band intensity and then purified using magnetic beads. A third and final PCR reaction on the purified pools was then used to add plate indexes and Illumina adapters: this reaction was performed in quadruplicate and the products from the four reactions were pooled together for purification. Sequencing was performed using the MiSeq Reagent Kit v3 on an Illumina MiSeq for 600 cycles (IBIS sequencing platform, Université Laval).

After sequencing, samples were demultiplexed using a custom python script with the reads being subdivided in four (plate barcode forward, row barcode, column barcode and plate barcode reverse). After demultiplexing, the forward and reverse reads were merged using the PANDA-Seq software^57^. Reads were then aligned to reference locus sequences using the Needle software from EMBOSS^58^. A custom script was then used to parse the alignments and extract genotype information for each read. The sequencing reads for the base editing deep sequencing experiment were deposited on the NCBI SRA as accession number PRJNA552472.

### Library transformation in yeast

Competent BY4741 (*MATa his3* Δ *1 leu2* Δ *0 met15* Δ *0 ura3* Δ *0*) cells were first transformed with the pKN1252 (p315-GalL-Target-AID) plasmid using a standard lithium acetate method. Transformants were selected by plating cells on SC-L. After 48 h of growth, multiples colonies were used to inoculate a starter liquid culture for competent cells preparation using the standard lithium acetate protocol^59^: a culture volume of 200 ml was used to generate enough competent cells for mass transformation. The large-scale library transformation was performed by combining 40 transformation reactions performed with 40 ul of competent cells and 5 ul of plasmid library (240 ng/ul) after the outgrowth stage and plating 100 ul aliquots on SC-UL: cells were then allowed to grow at 30°C for 48 h. A 1/1000 serial dilution of the cell recovery was plated in 5 replicates and used to calculate the number of transformants obtained. The total number of transformants reached 3.48 ×10^6^ CFU, corresponding to about 100X coverage of the plasmid pool.

### Target-AID mutagenesis and competition screening

The mutagenesis protocol is an upscaled version of our previously published method^23^ and is shown in Supplementary Figure 7. Transformants were scraped by spreading 5 ml sterile water on plates and then resuspending cells using a glass rake. All plates were pooled together in the same flask, and the OD of the yeast resuspension was measured using a Tecan Infinite F200 plate reader (Tecan, Switzerland). Pellets corresponding to about 6 x 10^8^ cells were washed twice with SC-UL without a carbon source and then used to inoculate a 100 ml SC-UL +2% glucose culture at 0.6 OD two times to generate replicates A and B. Cells were allowed to grow for 8 hours before 1 x 10^9^ cells were pelleted and used to inoculate a 100 ml SC-UL + 5% glycerol culture. After 24 hours, 5 x 10^8^ cells were pelleted and either put in SC-UL + 5% galactose for mutagenesis or SC-UL + 5% glucose for a mock induction control. Target-AID expression (from pKN1252) was induced for 12 hours before 1 x 10^8^ cells were pelleted and μ g/ml) co-selection culture in SC-ULR. After 16 hours of incubation, 5 x 10^7^ cells of each culture were used to inoculate 100 ml SC-UR, which was grown for 12 hours before 5 x 10^7^ cells were used to inoculate a final 100 ml SC-UR culture which was grown for another 12 hours. Cell pellets were washed with sterile water between each step, and all incubation occurred at 30°C with agitation. ∼2 x 10^7^ cells were taken for plasmid DNA extraction at the end of each mutagenesis and competition screening step.

### Yeast plasmid DNA extraction

Yeast plasmid DNA was extracted using the ChargeSwitch Plasmid Yeast Mini Kit (Invitrogen, California, USA) by following the manufacturer’s protocol with minor modifications: Zymolase 4000 U/ml (Zymo Research, California, USA) was used instead of lyticase, and cells were incubated for 1 hour at room temperature, one min at −80°C, and then incubated for another 15 minutes at room temperature before the lysis step. Plasmid DNA was eluted in 70 μ (10 mM Tris-HCl, pH 8.5) and stored at −20°C for use in library preparation.

### Next-generation library sequencing preparation

Libraries were prepared by using two PCR amplification steps, one to amplify the gRNA region of the pDSYCKO plasmid pool and the second to add sample barcodes as well as the Illumina p5 and p7 sequences^60^. Oligonucleotides for library preparation are shown in the first part of the oligonucleotide table. Reaction conditions for the first PCR were as follows:

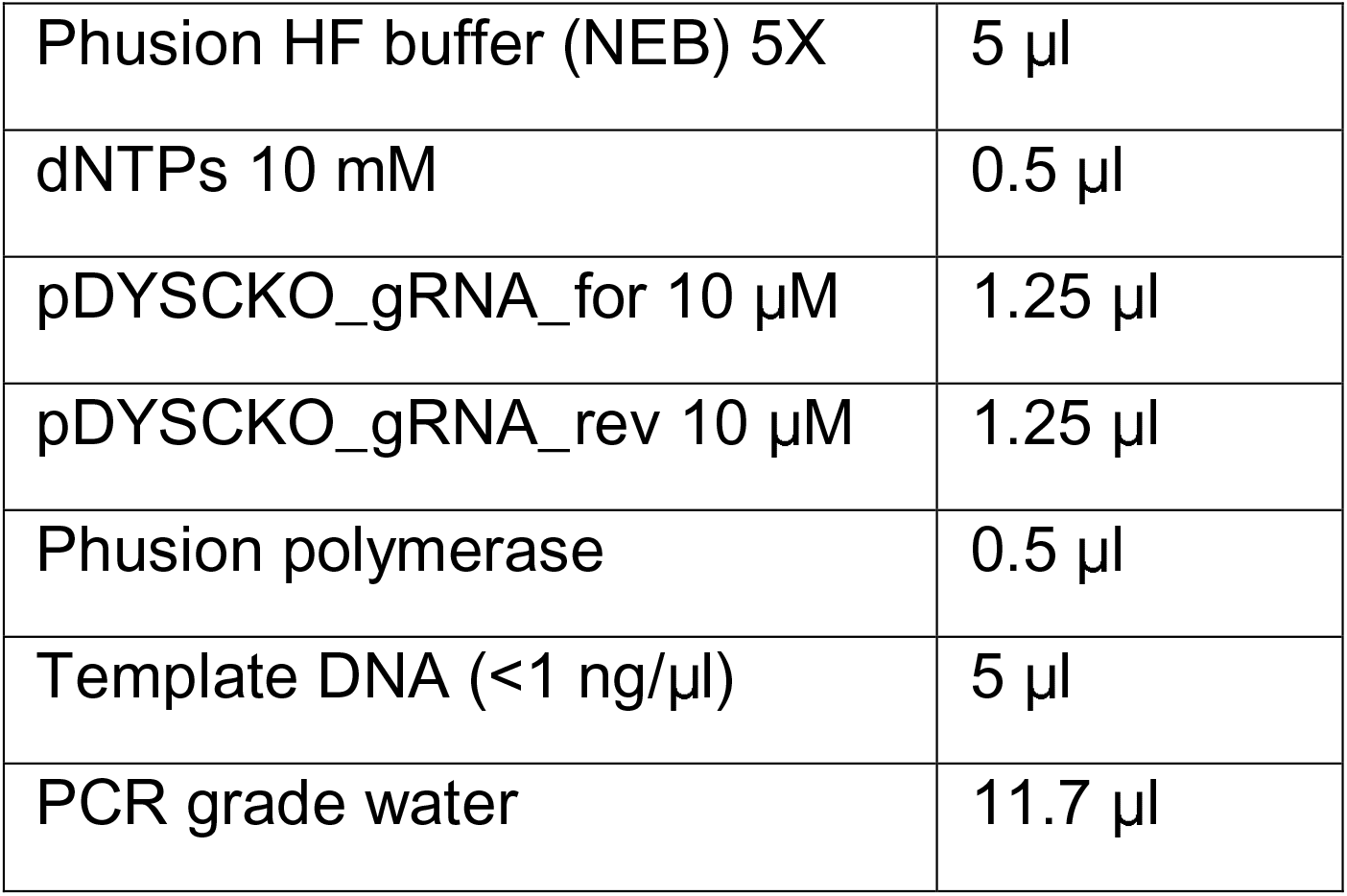

Thermocycler protocol:

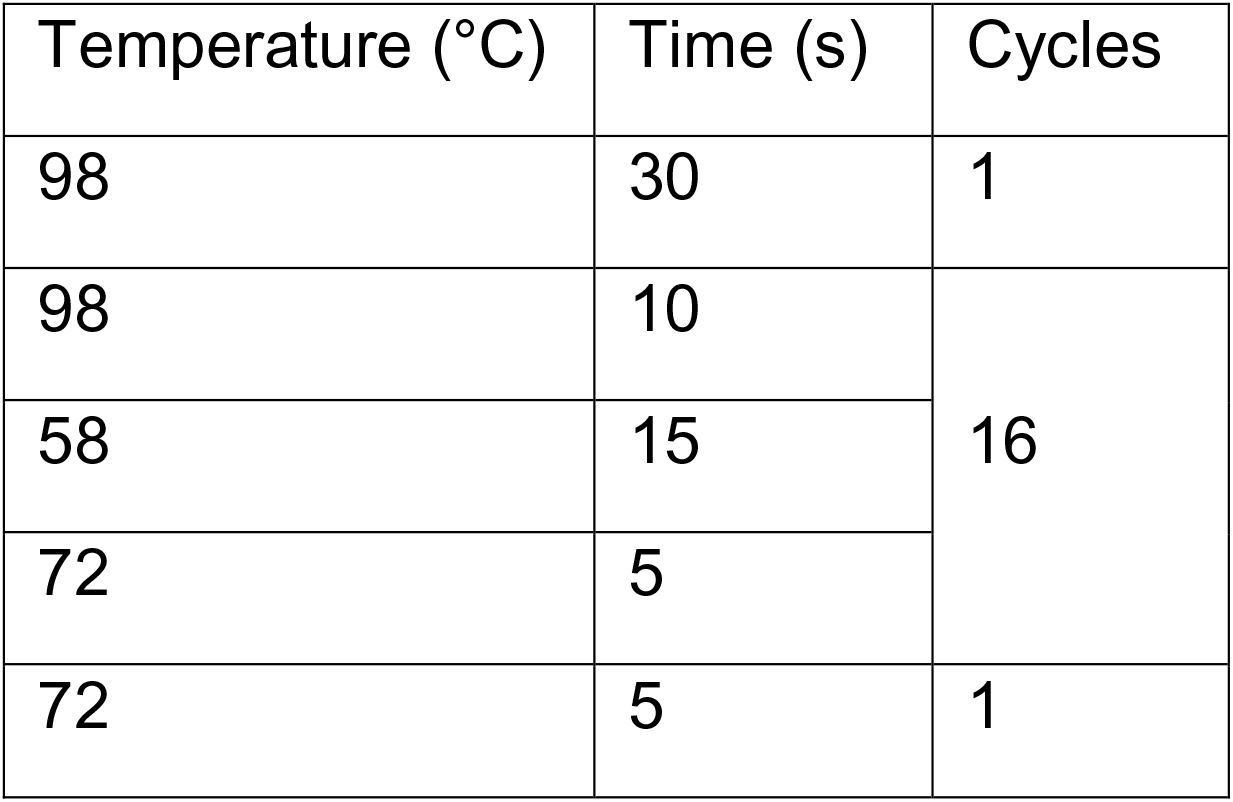

The resulting product was verified on a 2% agarose gel colored with Midori Green Advance (Nippon Genetics, Japan) and then gel-extracted and purified using the FastGene Gel/PCR Extraction Kit (Nippon Genetics, Japan). The purified products were used as the template for the second PCR reaction, with the following conditions:

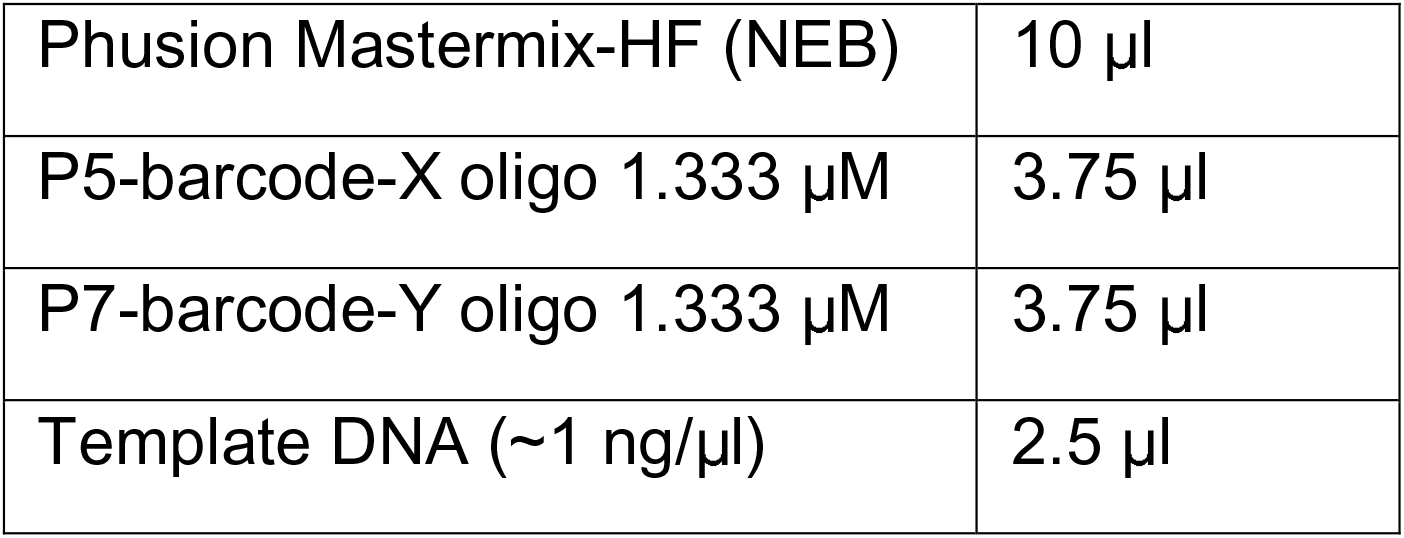

Thermocycler protocol:

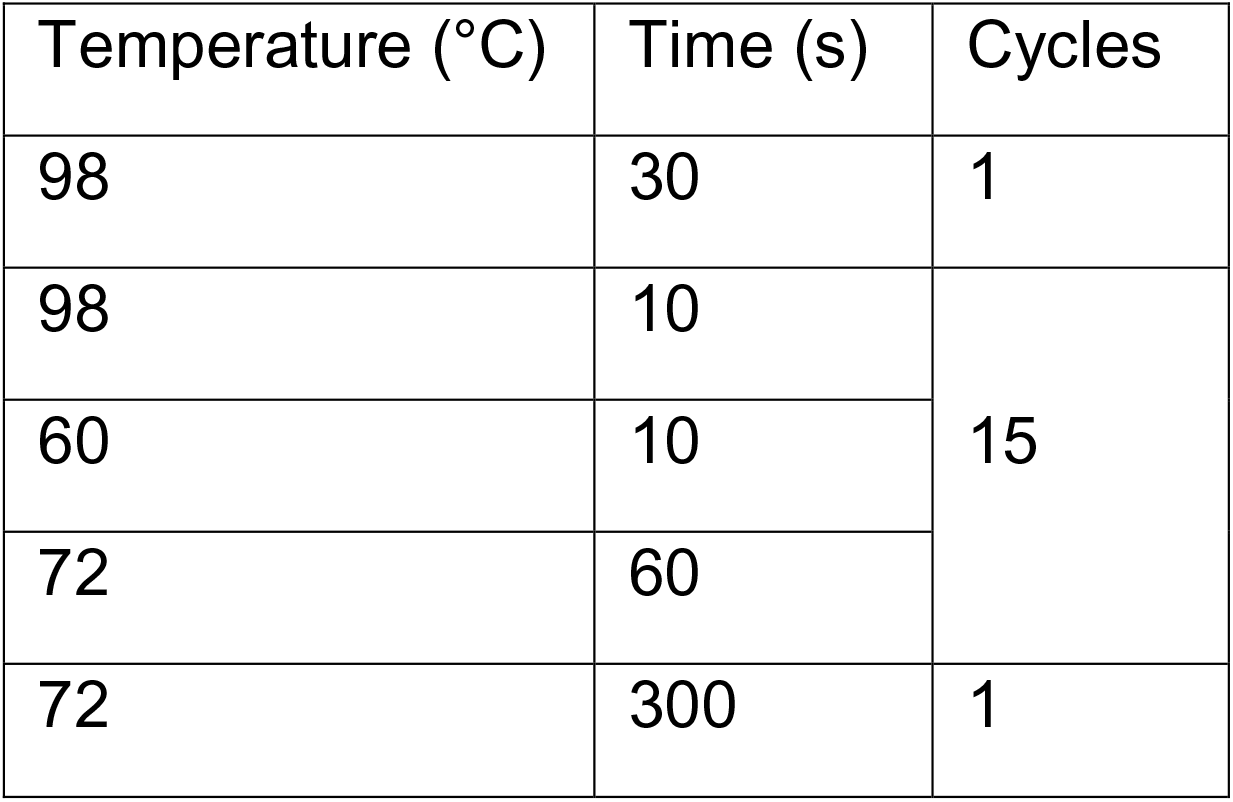

PCR products were verified on a 2% agarose gel colored with Midori Green Advance (Nippon Genetics, Japan) and then gel-extracted and purified using the FastGene Gel/PCR Extraction Kit (Nippon Genetics, Japan). Library quality control and quantification were performed using the KAPA Library Quantification Kit for Illumina platforms (Kapa Biosystems, Massachusetts, USA) following the manufacturer’s instructions. Libraries were then run on a single lane on HiSeq 2500 (Illumina, California, USA) with paired-end 150 bp in fast mode.

### Large-scale screen sequencing data analysis

The custom Python scripts used to analyze the are available on github (https://github.com/landrylaboratory), and packages and software used are presented in Supplementary table 9. Raw sequencing files have been deposited on the NCBI SRA, accession number PRJNA552472. Briefly, reads were separated into three subsequences for alignment: the P5 barcode, the gRNA, and the P7 barcode. Each of these was aligned using Bowtie ^50^ to an artificial reference genome containing either the barcodes or gRNA sequences flanked by the common amplicon sequences. The gRNA sequences are aligned both with 0 or 1 mismatch allowed, and misalignment position and type were stored. Information on barcode and gRNA alignment for each read was stored and combined to generate a barcode count per library table, a list of mismatches in alignments for each gRNA in each library, as well as mismatch types and counts for the same gRNA across all libraries.

gRNAs absent from more than half of the libraries (4446 out of 39,989) were removed from the analysis before gRNA abundance calculations.

### Detecting mutations with high fitness effects

Barcode sequencing competition experiments use DNA barcodes to measure the relative abundance of many different subpopulations of cells grown in the same pool (Robinson *et al.* 2014). Since each gRNA is linked to its possible mutagenesis outcomes, we can use relative gRNA abundance to detect mutations with significant fitness effects. To do so, the log_2_ of the relative abundance of a barcode after mutagenesis is compared with its abundance at the end of the screen:

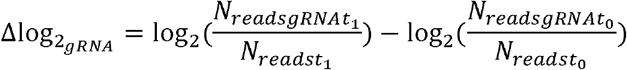

For each gRNA, the measured fitness effect is the product of the effect of the mutational outcomes on growth and of the mutation rate within the cell subpopulation bearing this particular gRNA. Relative counts will also vary stochastically because of variation in sequencing coverage depending on the time point and replicate. To reduce the impact of these effects, a minimal read count at the end of the galactose induction step was used to filter out low abundance gRNAs. We found a minimal read threshold of n=54 provided a good tradeoff between the number of gRNAs eligible for analysis and inter-replicate correlation.

To obtain a reference distribution of abundance variation for gRNAs, we fit a normal distribution Δlog2 z-score distribution of gRNAs between the start of the experiment (The glucose timepoint in Supplementary Figure 9) and the end of the mock glucose induction time point (n=7875 values). The mock induction recapitulates the galactose induction time point, but using glucose as a sole carbon source so that Target-AID is not expressed. Using this reference distribution, we calculated a z-score for each gRNA during the competition experiment independently for both replicates. We then averaged z-scores between replicates. We set a significance threshold such as that all gRNAs at z-scores for which the estimated False Discovery Rate ∼5% and the False positive Rate ∼0.2% are considered GNEs (Supplementary Figure 9 B and C).

### Complementation assays

Experiments were performed in heterozygous deletion mutants from the YKO project heterozygous deletion strain set (Dharmacon, Colorado, USA). For each gene, a single colony streaked from the glycerol stock was used to prepare competent cells using the previously described lithium acetate protocol^59^. To generate mutant alleles of the genes of interest, we performed site-directed mutagenesis on the appropriate MoBY collection plasmid ^27^. These centromeric plasmids encode the yeast gene of interest under the control of their native promoters and terminators. Mutagenesis reactions were performed with the following reaction setup:

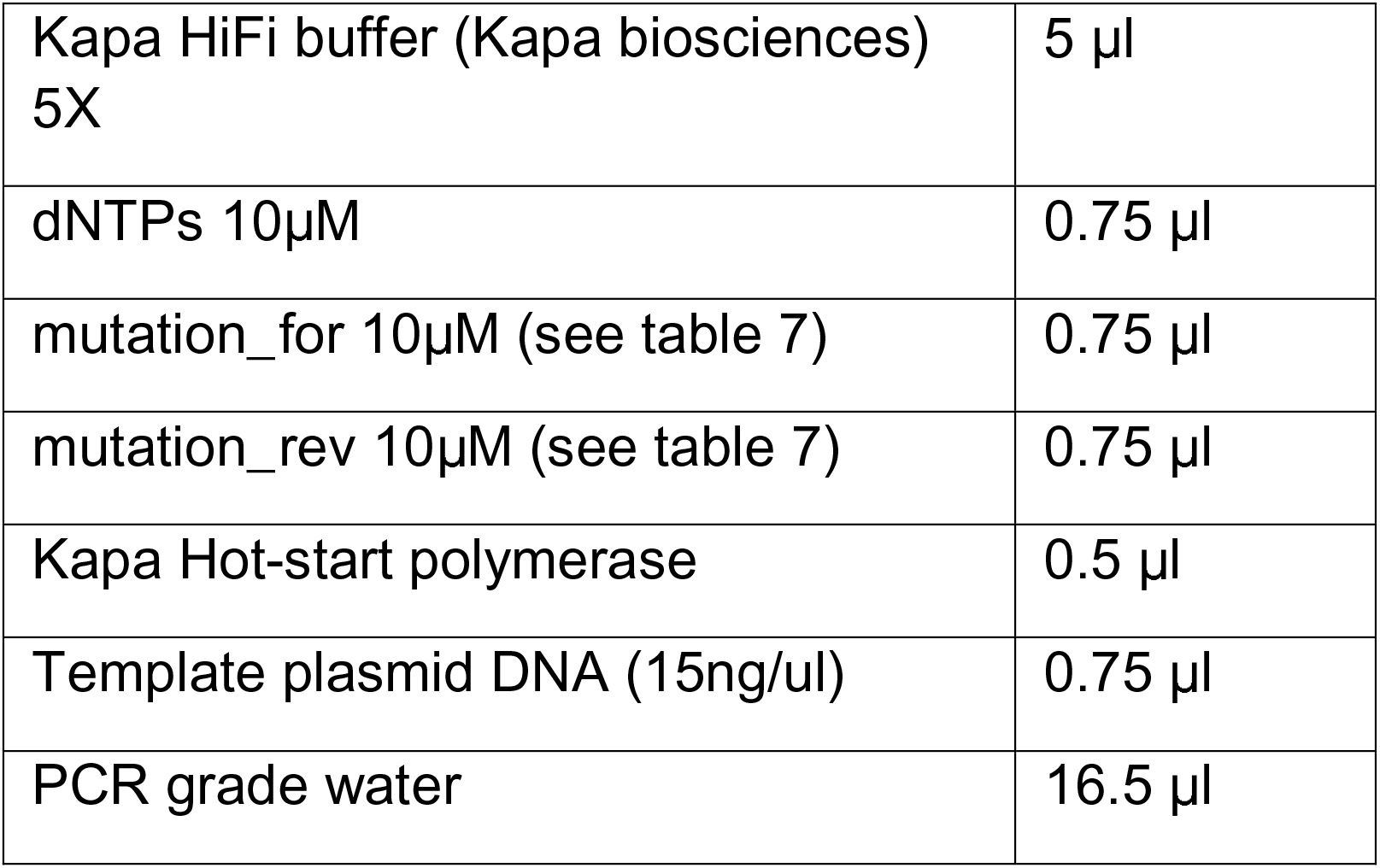

Thermocycler protocol:

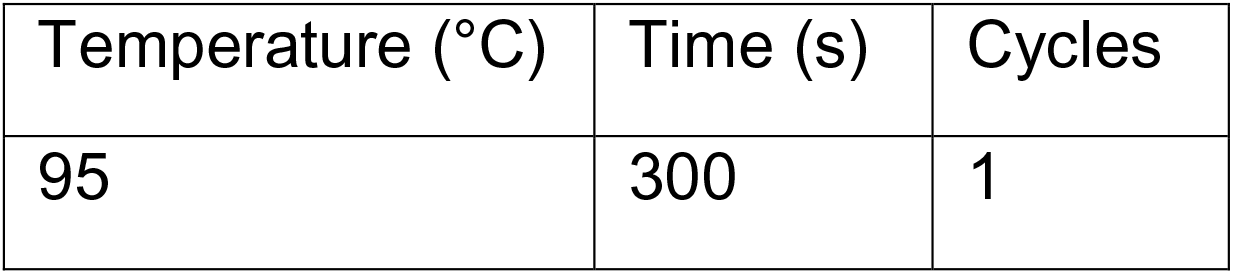

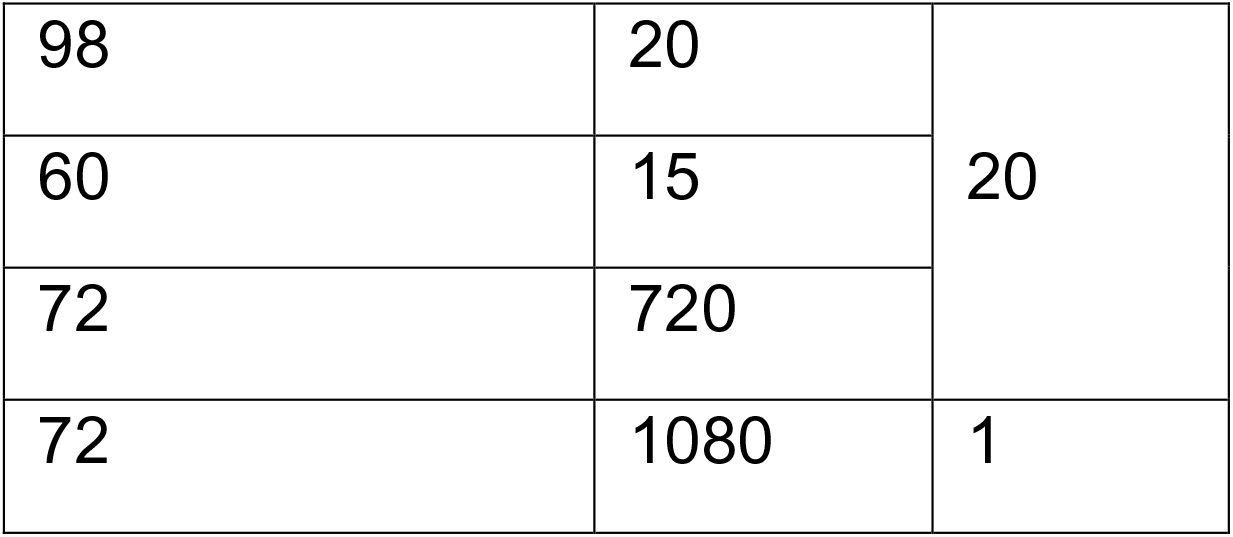

After amplification, the mutagenesis product was digested with DpnI for 2 hours at 37°C and 5 ul was transformed in *E. coli* strain BW23474 (F-, Δ *(argF-lac)169*, Δ*uidA4::pir-116*, *recA1*, *rpoS396(Am)*, *endA9(del-ins)::FRT*, *rph-1*, *hsdR514*, *rob-1*, *creC510*)^61^. Transformants were plated on 2YT+Kan+Chlo and grown at 37°C overnight. Plasmid DNA was then isolated from clones and sent for Sanger sequencing (CHUL sequencing platform, Université Laval, Québec City, Canada) to confirm mutagenesis success.

Competent cells of target genes were transformed with the appropriate mutant plasmids as well a the original plasmid bearing the wild-type gene and the empty vector^62^, and transformants were selected by plating on SC-U (MSG). Multiple independent colonies per transformation were then put on sporulation media until sporulation could be confirmed by microscopy. For tetrad dissection, cells were resuspended in 100ul 20T zymolyase (200mg/ml dilution in water) and incubated for 20 minutes at room temperature. Cells were then centrifuged and resuspended in 50ul 1M sorbitol before being streaked on a level YPD plate. All dissections were performed using a Singer SporePlay microscope (Singer Instruments, UK). Plate pictures were taken after five days incubation at room temperature except for the RAP1 plasmid complementation test for which the picture was taken after three days. Pictures are shown in Supplementary Image File 1.

### Strain construction for confirmations in *RAP1*

Because the MoBY collection plasmid for RAP1 cannot fully complement the gene deletion (Supplementary image file 1), we instead performed confirmations by engineering mutations a diploid strain to create heterozygous mutants. *RAP1* was first tagged with a modified version of fragment DHFR F[1,2] (the first half) of the mDHFR enzyme^63^. The mDHFR[1,2]-FLAG cassette was amplified using gene-specific primers and previously described reaction parameters^63^. Cells were transformed with the cassette using the previously described transformation protocol and were plated on YPD+Nourseothricine (YPD+Nat in Media table). Positive clones were identified by colony PCR and successful fragment fusion was confirmed by Sanger sequencing (CHUL sequencing platform). We then mated the confirmed clones with strain Y8205 (*Mat* α *can1::STE2pr-his5 lyp1::STE3prLEU2* Δ*ura3* Δ*his3* Δ*leu2*, Kindly gifted by Charlie Boone) by inoculating a 4ml YPD culture with overnight starter cultures of both strains and letting the culture grow overnight. Cells were then streaked on YPD+Nat and diploid cells were identified by colony PCR using mating type diagnosis primers^64^.

To create heterozygous deletion mutants of the target gene, we amplified a modified version of the *URA3* cassettes that could then be targeted with the CRISPR-Cas9 system to integrate our mutations of interest using homologous recombination at the target locus. The oligonucleotides we used differ from those commonly used in that they amplify the cassette without the two LoxP sites present at both ends. We found it necessary to remove those sites as one common mutational outcome after introducing a double-stranded break in the *URA3* cassette was inter-LoxP site recombination without the integration of donor DNA at the target locus. These modified cassettes recombine with DNA upstream the target gene on one end and the mDHFR F[1,2] fusion on the other, ensuring that the heterozygous deletion is always performed at the locus that is already tagged. Cassettes were transformed using the standard lithium acetate method, and cells were plated on SC-U (MSG) selective media. Heterozygous deletion mutants were then confirmed by colony PCR.

### CRISPR-Cas9 mediated Knock-in of targeted mutations

Mutant alleles of target genes were amplified in two fragments using template DNA from the haploid tagged strain (See Supplementary figure 14). The two fragments bearing mutations were then fused together by a second PCR round to form the final donor DNA. This DNA was then co-transformed with a plasmid bearing Cas9 and a gRNA targeting the URA3 cassette for HDR mediated editing using a standard protocol^65^. Clones were then screened by PCR to verify donor DNA and mutation integration at the target locus. The targeted region of *RAP1* was then Sanger sequenced (CHUL sequencing platform, Univesité Laval, Québec City, Canada) to confirm the presence of the mutation of interest. Heterozygous mutants were sporulated on solid media until sporulation could be confirmed by microscopy using the same protocol previously described. The plates were then replica plated on YPD+Nat media, and the pictures were taken after five days at room temperature (Supplementary Image File 2).

### Evolutionary rate measurements and protein variant abundance

Evolutionary rates were calculated using the Rate4site software^66^ using multiple sequence alignments and phylogenies from PhylomeDB V4^67^ as input and using the raw calculated rates as output. Variant data was compiled using data from the 1002 Yeast Genome Project (http://1002genomes.u-strasbg.fr/files/allReferenceGenesWithSNPsAndIndelsInferred.tar.gz). Strain-specific protein coding sequence were aligned to the S288c sequence using Fastx36^68^ with the following parameters: `fastx36 -p -s -VT10 -T 6 -m 10 -n -3 querymultifasta.fasta ref_orf.db 12\> fasta_out`. Alignments were then parsed with a custom Python script to identify variants. Variant abundance was measured as the number of strains in the dataset in which a specific variant was found. If the coding sequence contained ambiguous nucleotides (ex.: R or Y), separate coding sequences were generated for each possibility and each possible variant was considered as a separate occurrence.

### Analysis of the properties of stop codon generating gRNAs

To analyse the sequence and target properties of gRNA inducing the creation of stop codons, data from multiple sources was compiled. For each target gene, length and chromosomal strand was obtained from the Saccharomyces Genome Database using the Yeastmine query interface^69^. Distance to centromere was obtained by calculating the minimal distance between the start of the gene and one extremity of the centromere coordinates. RNA:DNA duplex melting temperature of gRNA sequence with target genomic DNA was calculated using the MeltingTemp module from Biopython^70^, which uses values taken from Sugimoto et al^71^. Correlation between gRNA/DNA duplex melting temperatures was assessed using Spearman’s rank correlation.

### Variant effect prediction resources analysis and GO enrichment

All prediction data except the Envision scores were extracted from the aggregated data of the Mutfunc database^31^. Precomputed values were downloaded directly from the FTP server (http://ftp.ebi.ac.uk/pub/databases/mutfunc/mutfunc_v1/yeast/). This database includes precomputed SIFT scores for 5498 yeast proteins, as well as predicted variant ddG values based on protein structure (n=1057), homology models (n=1703) and protein-protein interaction interfaces (n=1109). Mutations with ΔΔ G>1 considered destabilizing.

Precomputed values from Envision^2^ were downloaded directly from the database website (https://envision.gs.washington.edu/shiny/envision_new/, file yeast_predicted_2017-03-12.csv). This file contained 34857830 mutation effect predictions spread across 4011 genes. The distribution of Envision scores for the genes targeted in the experiment that are included in the database are shown in Supplementary Figure 12.

We downloaded the Uniprot database for yeast genes (query: uniprot-proteome_UP000002311) with annotations covering the following properties: Metal binding, Nucleotide binding, Site, DNA binding, Calcium binding, Binding site, Active site, Motif. We found that 6295 gRNAs targeted genes which have annotations in Uniprot, of which 519 were GNEs (ratioAll=0.0749). Statistical enrichments were calculated using this set of gRNAs as the reference population. Gene enrichments were performed using the PANTHER gene list analysis tool^72^. The list of genes for which 2 or more GNEs were detected was tested for enrichment against all genes targeted by the library using Fisher’s exact test and False Discovery Rate calculations. The Gene Ontology datasets used were: GO molecular function complete, GO biological process complete, and GO cellular component complete.

## Supporting information

Source image File 1

Source image File 2

## Data Availability

All raw sequencing data has been deposited on the NCBI as accession number PRJNA552472. The gRNA screen scores, predicted mutation outcomes, mutation effect predictors scores, as well as other relevant annotations are presented in Supplementary Dataset 1. Source image files for the tetrad dissections are presented as Supplementary Image 1 and 2.

## Code Availability

The custom Python scripts used to analyze the are available on github (https://github.com/landrylaboratory), and packages and software us ed are presented in Supplementary table 9.

## Acknowledgments

This work was supported by the Canadian Institutes of Health Research Foundation grant 387697 to CR.L., as well as project grants 364920, 384483, a Frederick Banting and Charles Best graduate scholarship and a Vanier graduate scholarship to P.C.D, by Université Laval via an André Darveau Fellowship to P.C.D., the Fonds Québécois de Recherche en Santé via a Master’s training award to P.C.D. and the Japan Society for the Promotion of Science grant numbers S15734 and S17161 to C.R.L. and N.Y. The authors thank Mathieu Hénault, Johan Hallin, and Dan Yamamoto Evans for comments on the manuscript, as well as Maria Isabel Acosta Lopez for assistance during the strain construction process.

## Author contributions

PCD, AKD, NY and CRL designed research. PCD and AKD performed experiments. PCD and MS generated NGS sequencing data. All data analysis was performed by PCD with input from CRL. PCD and CRL wrote the manuscript with input from all authors.

## Conflict of interest

None to declare

## Notes

#### Summary of Updates

Change to figure 3, methods, and supplementary materials

