## Supplementary material for "Perturbing proteomes at single residue resolution using base editing": Source image File 1

### RAP1

MoBY WT MoBY empty

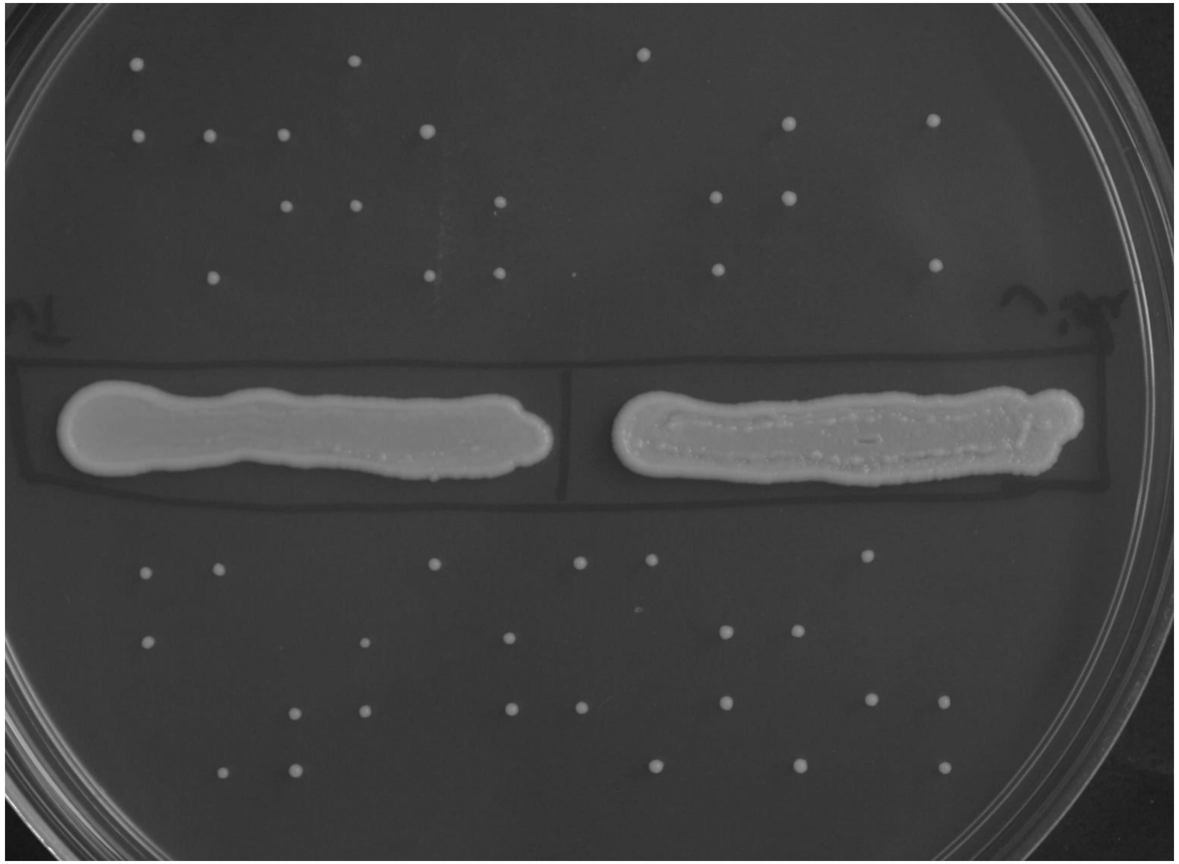

### GLN4

MoBY WT

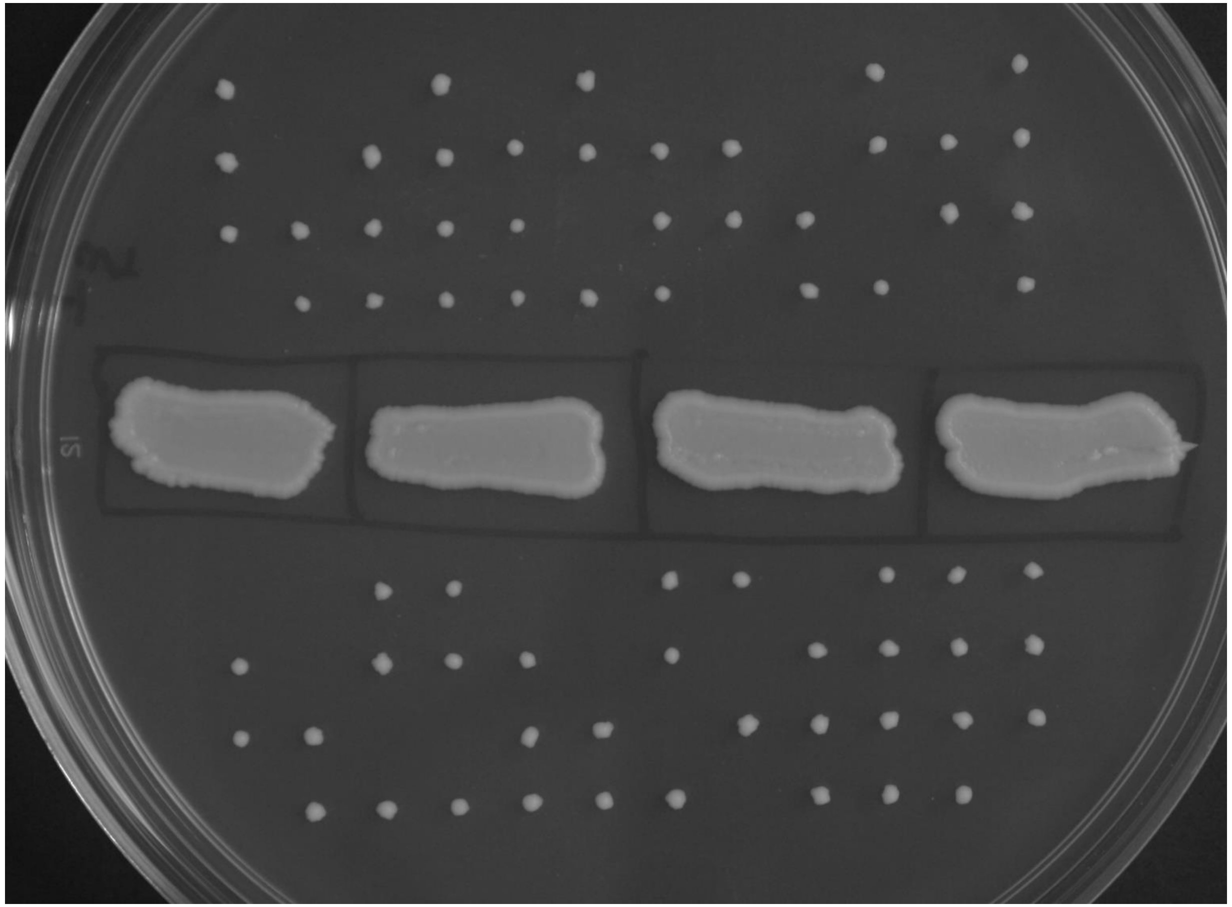

MoBY empty

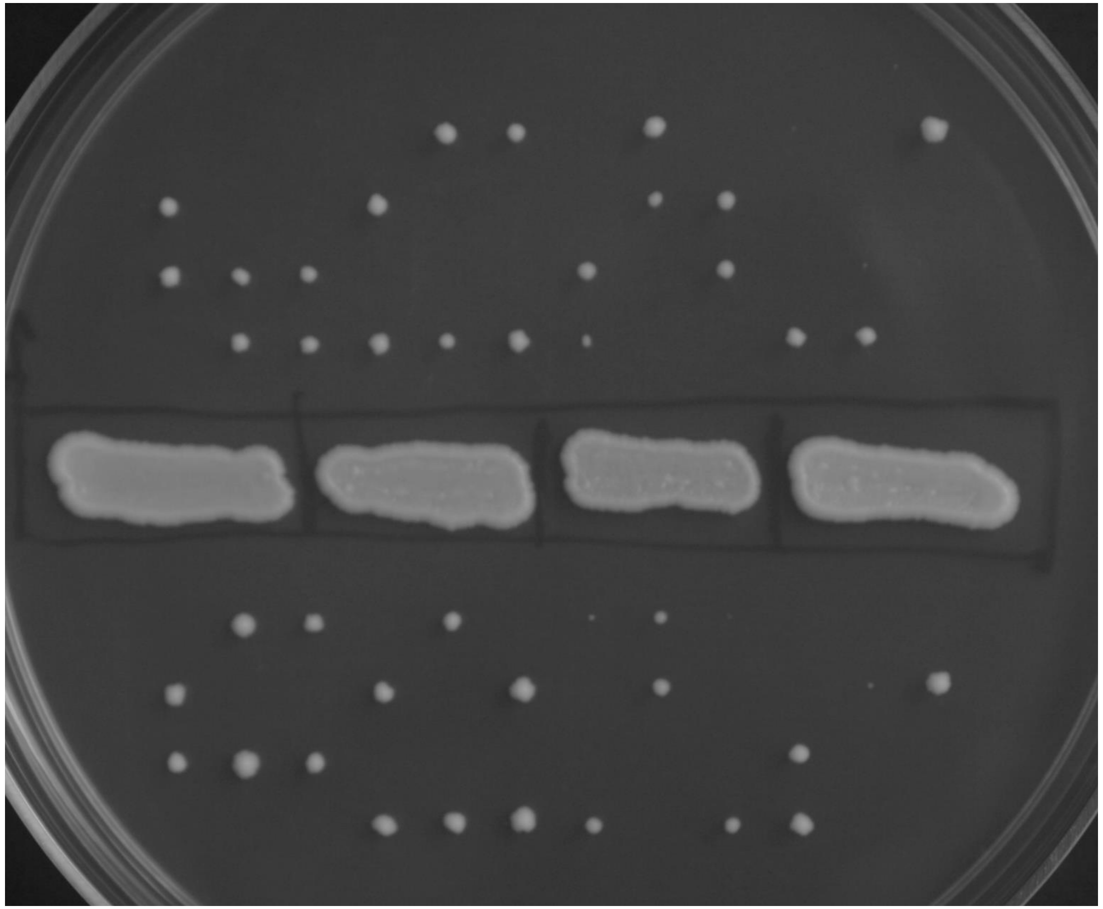

### GLN4

G267R

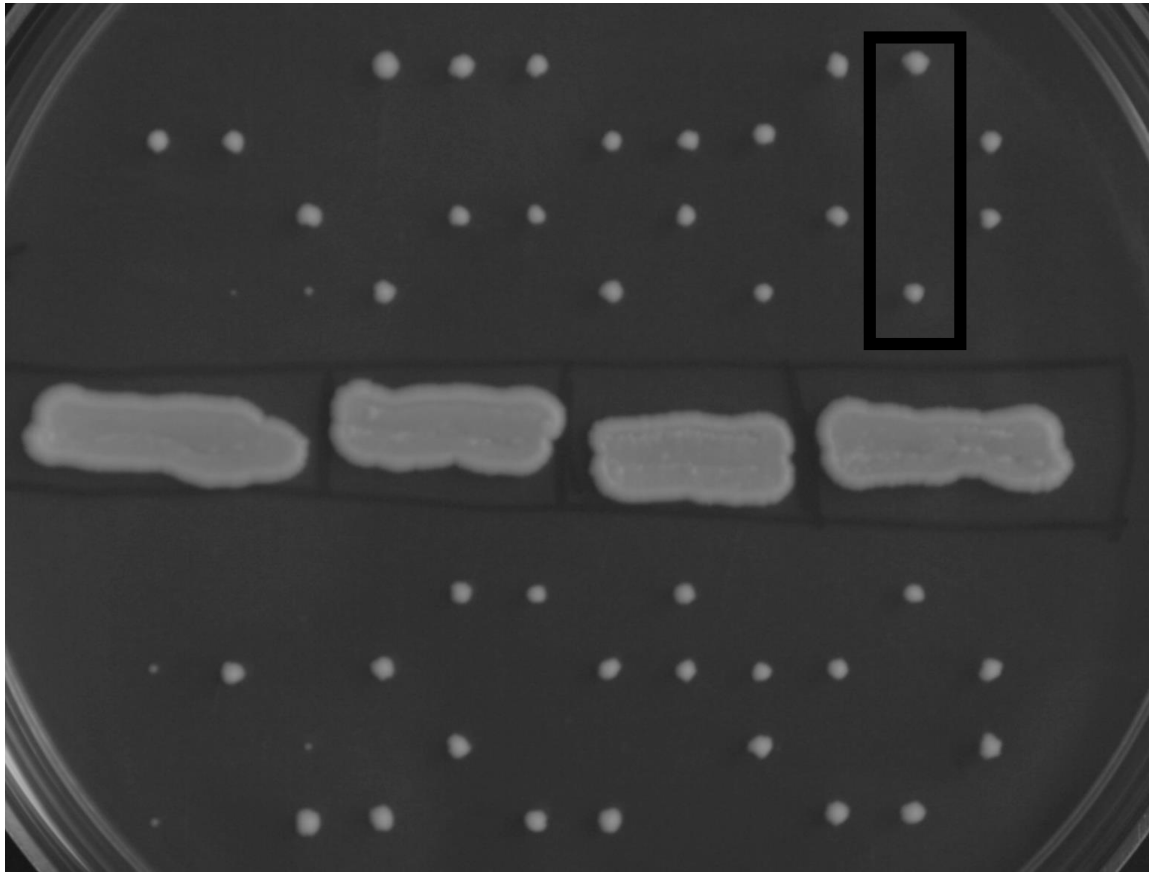

G267S

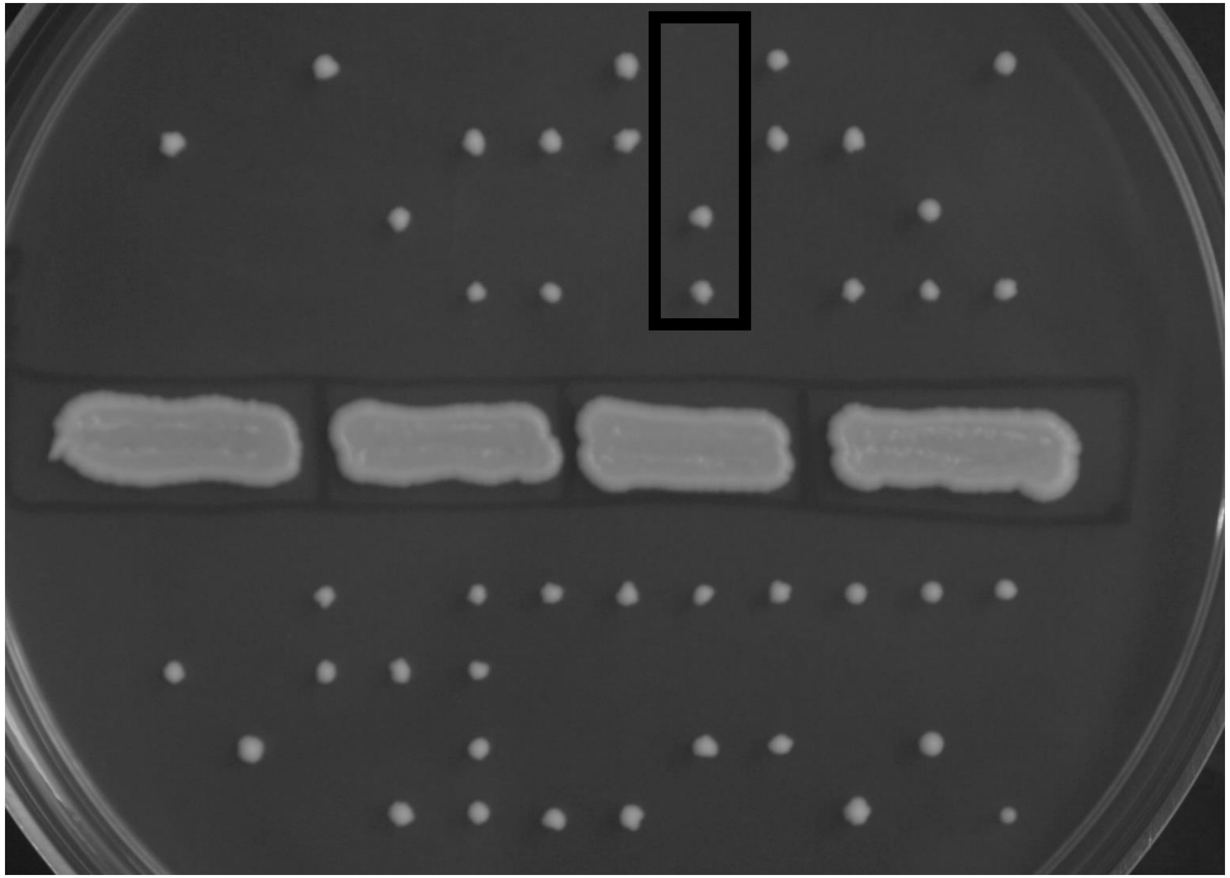

### GLN4

D291E D291D

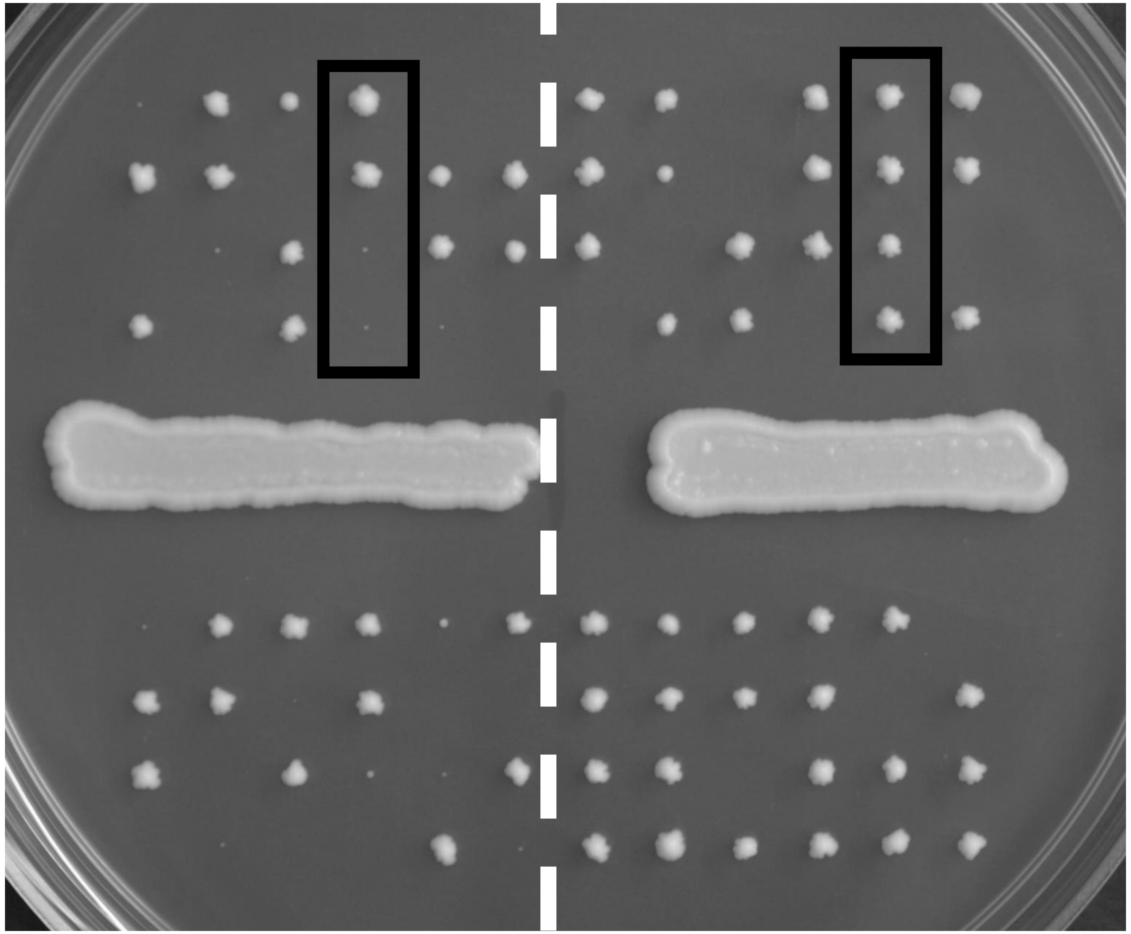

R402T R402K

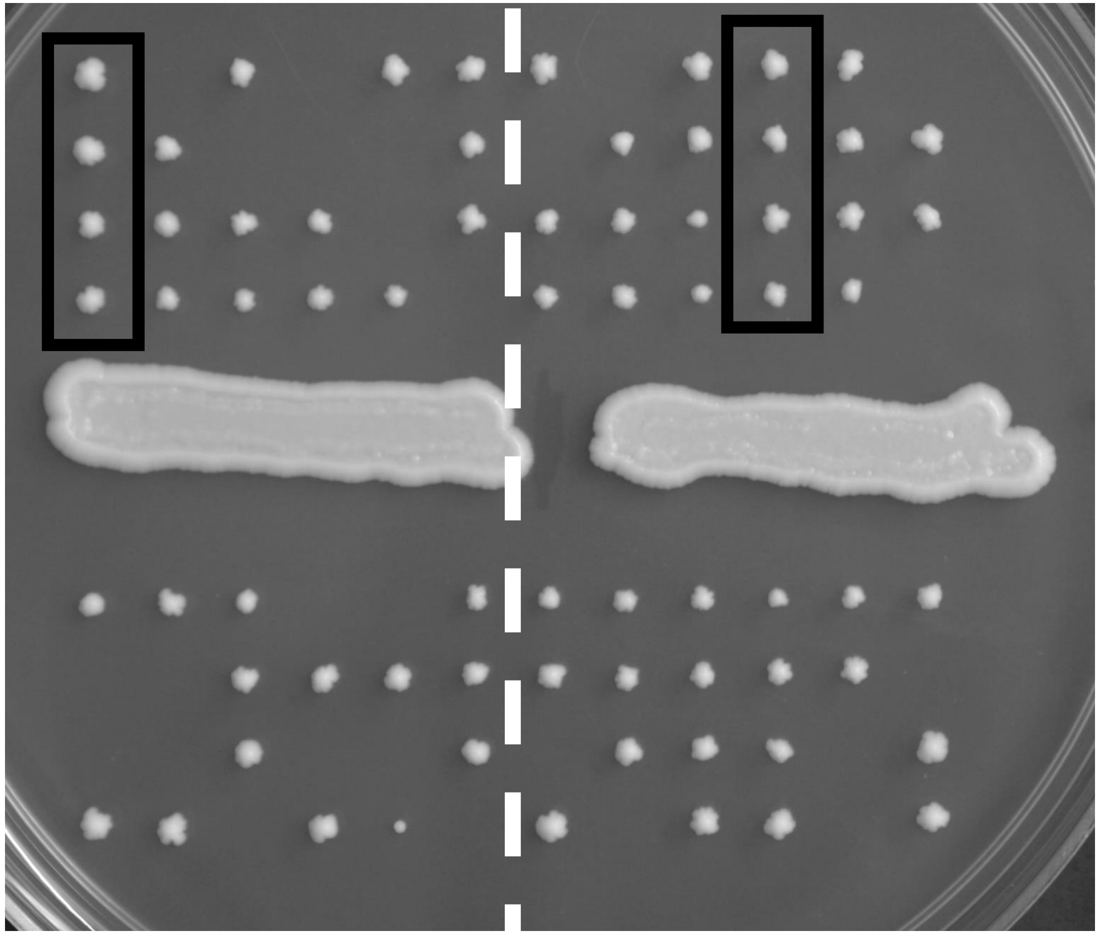

### GLN4

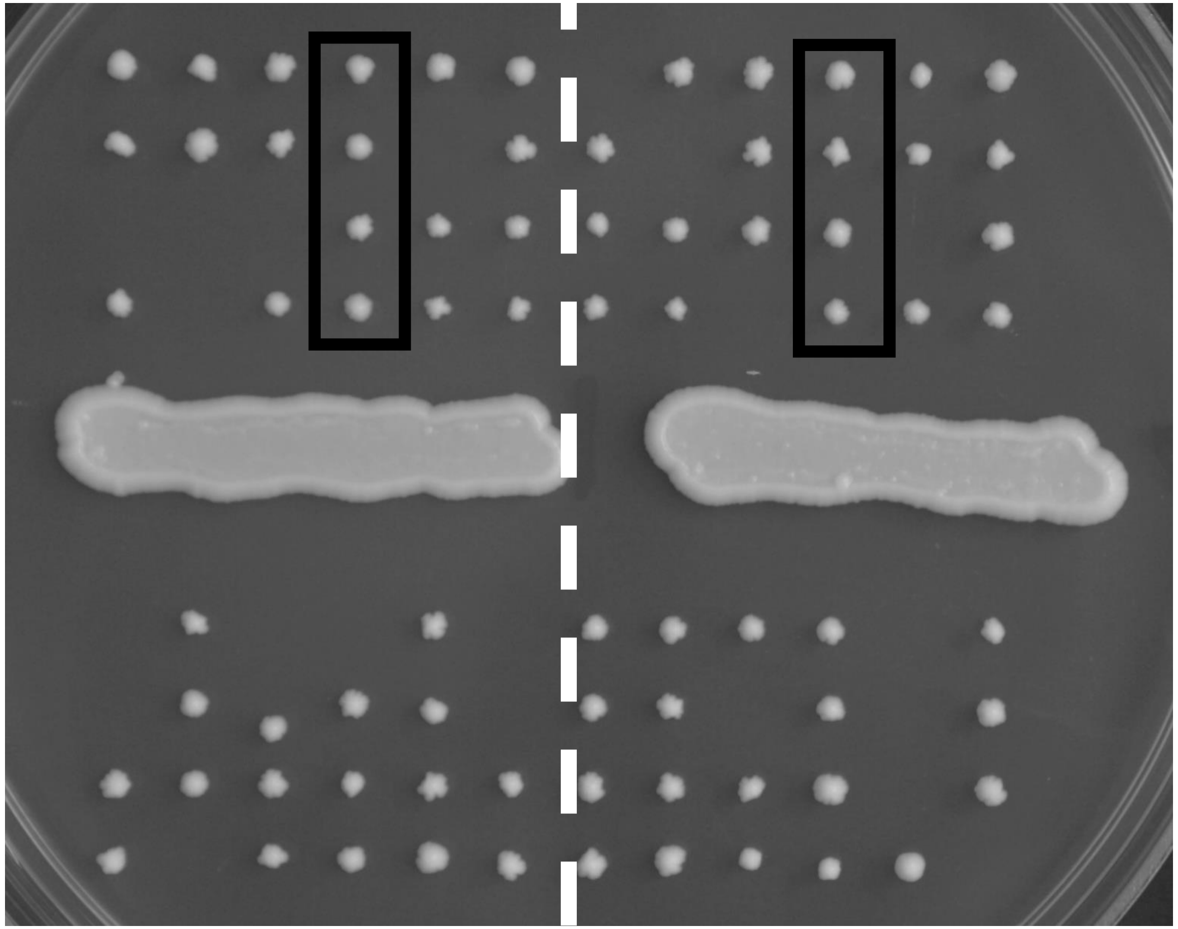


T548R

NA

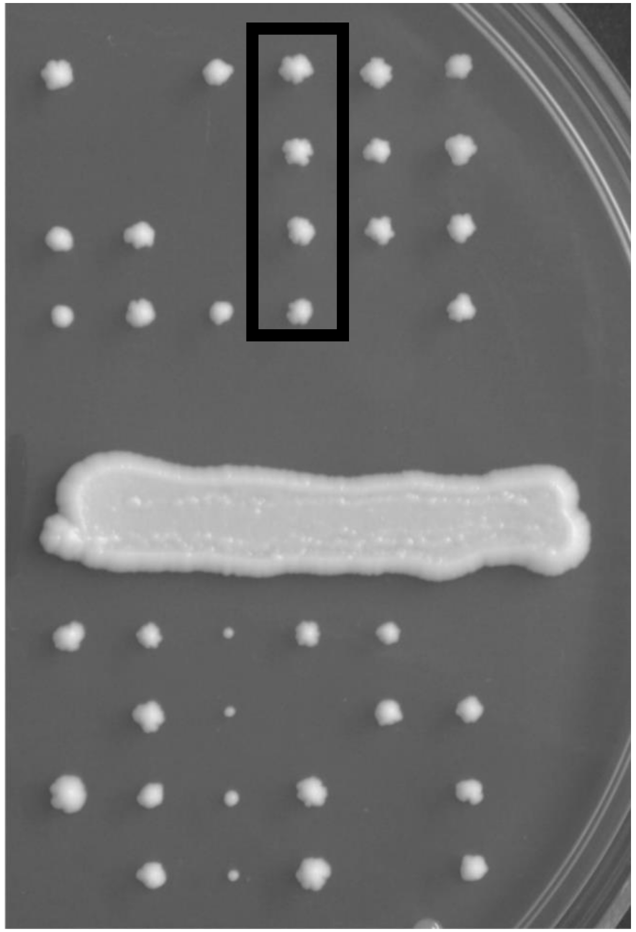

### GLN4

R569T

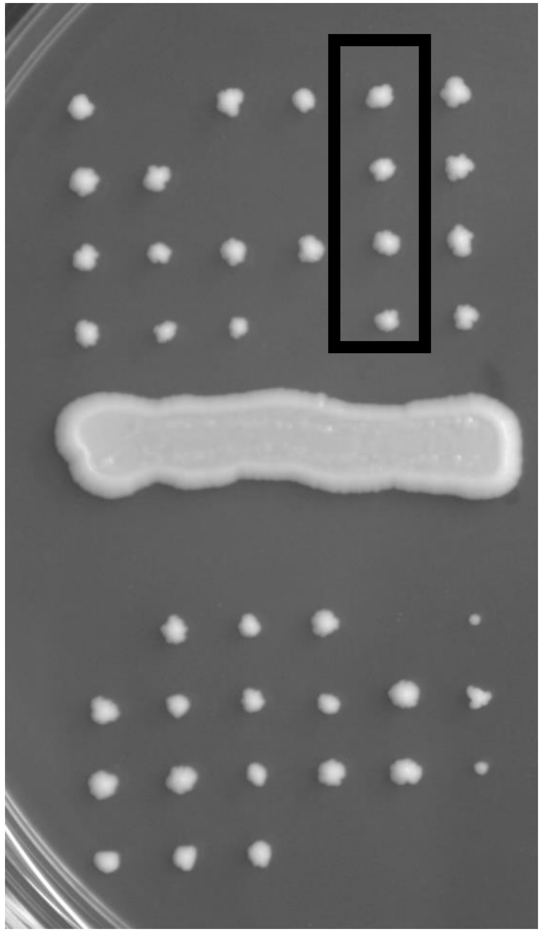

R569K

NA

E301Q

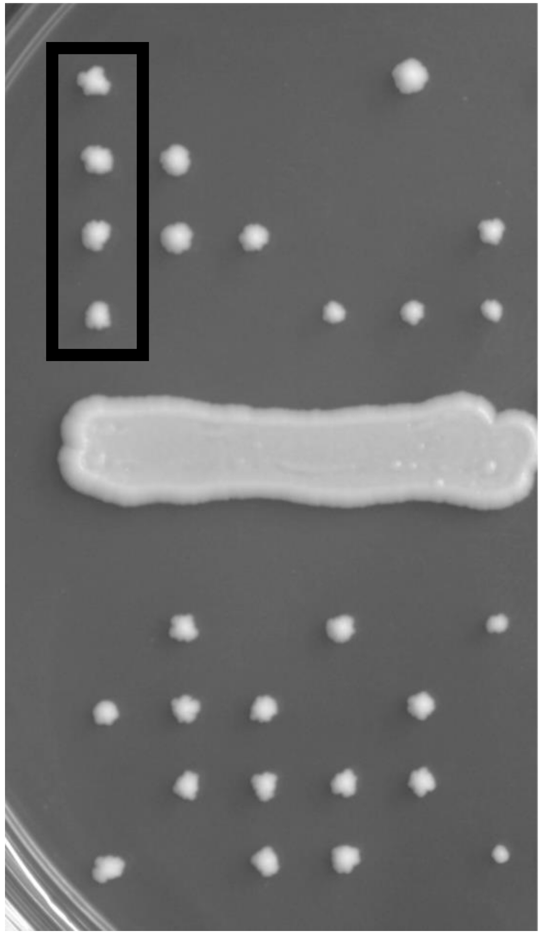

E301R

NA

### GLN4

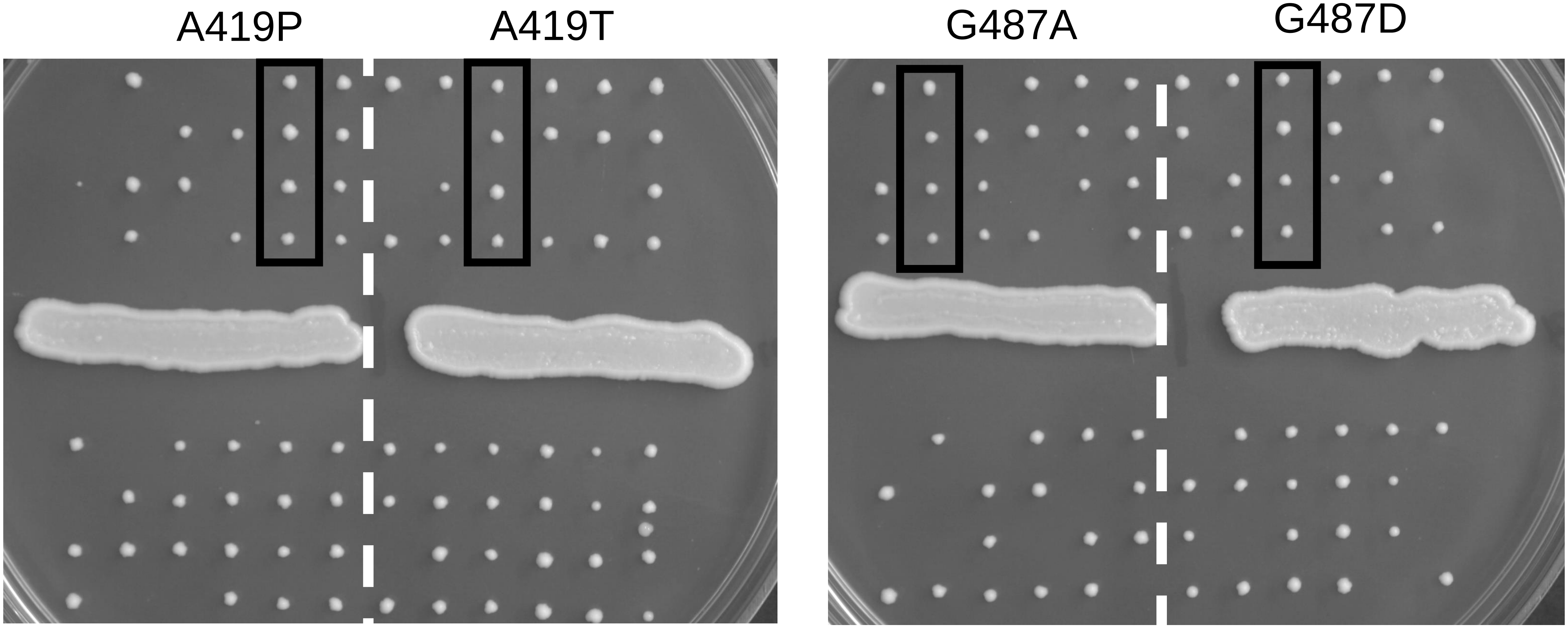

### GLN4

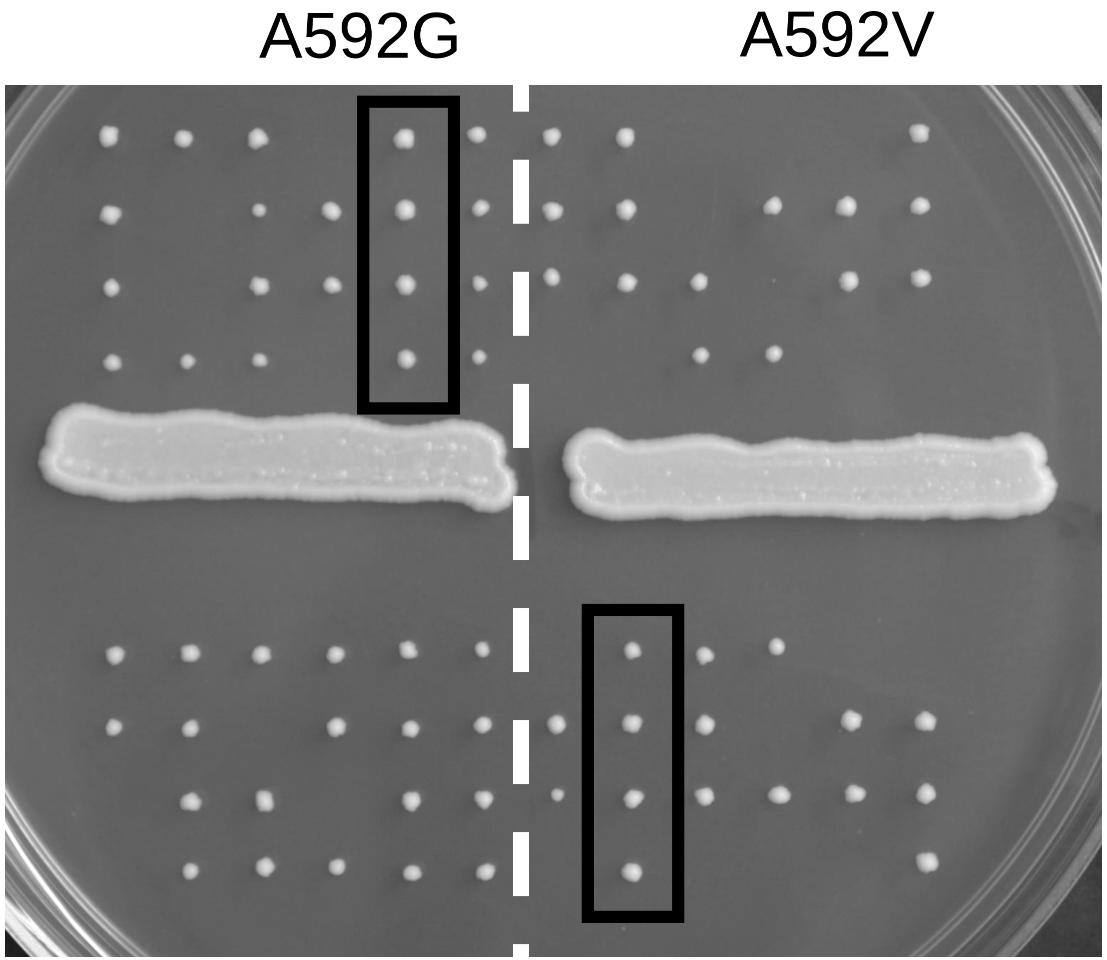
