## Supplementary figures and images for "Perturbing proteomes at single residue resolution using base editing"

### Source image File 2

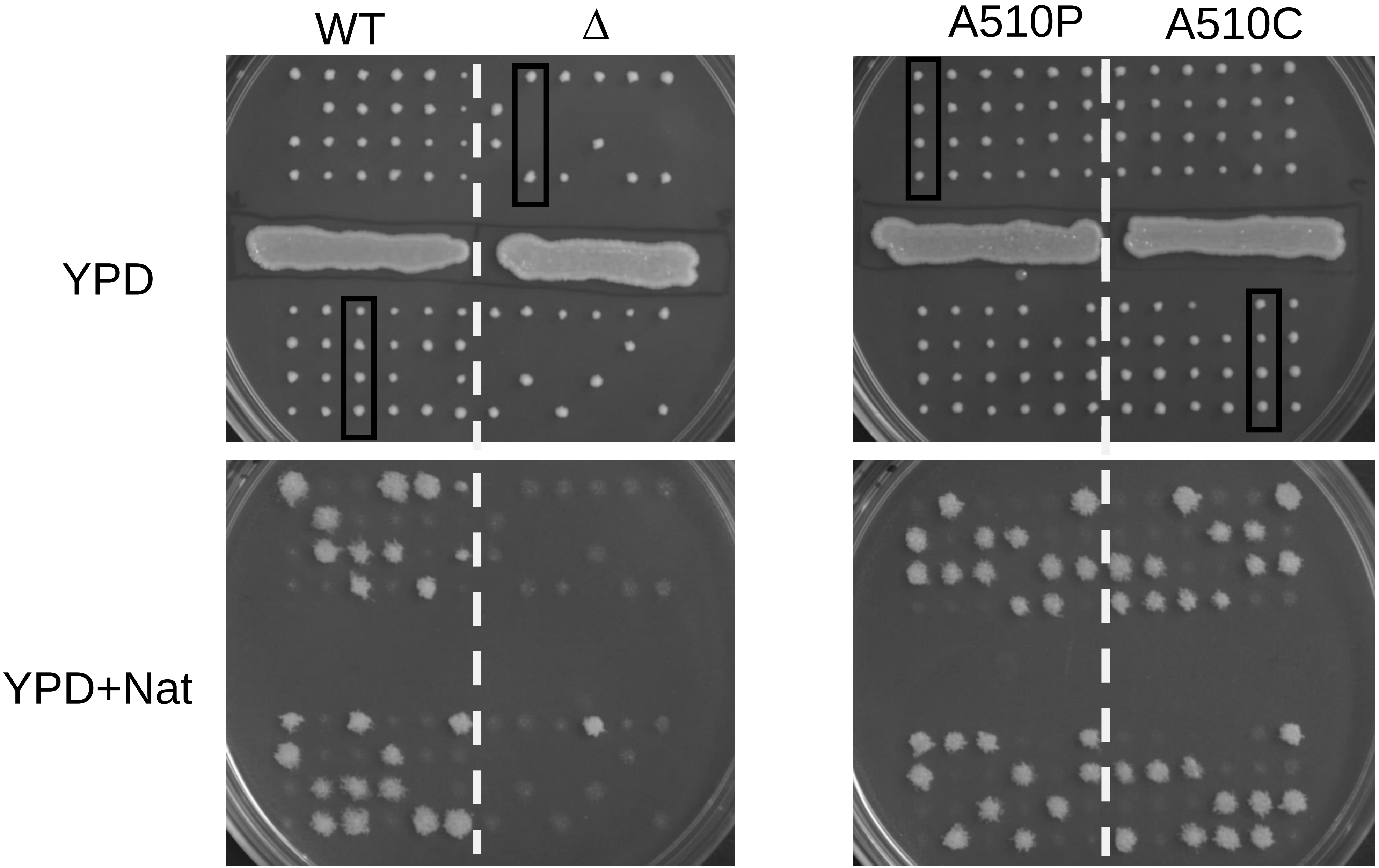

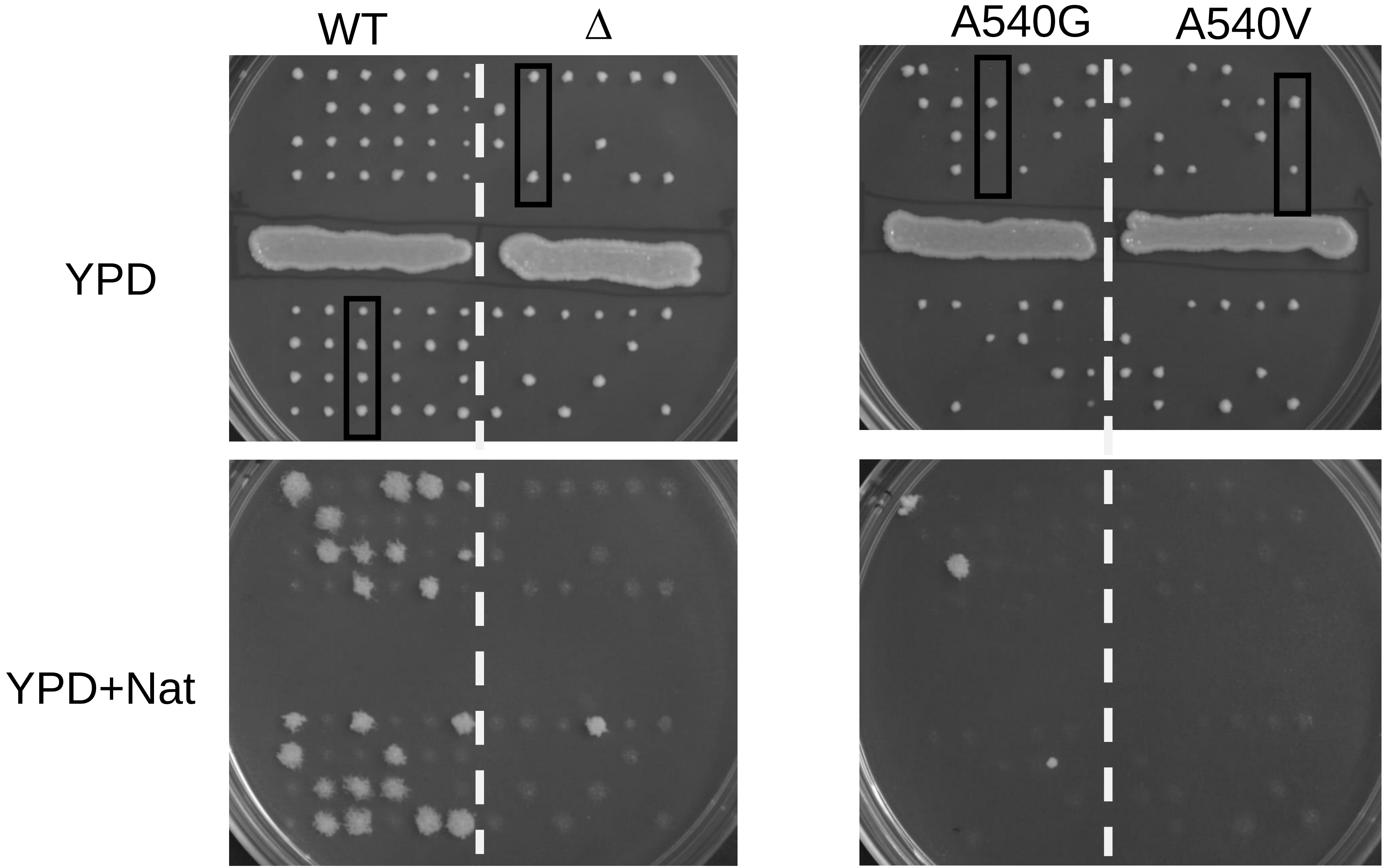

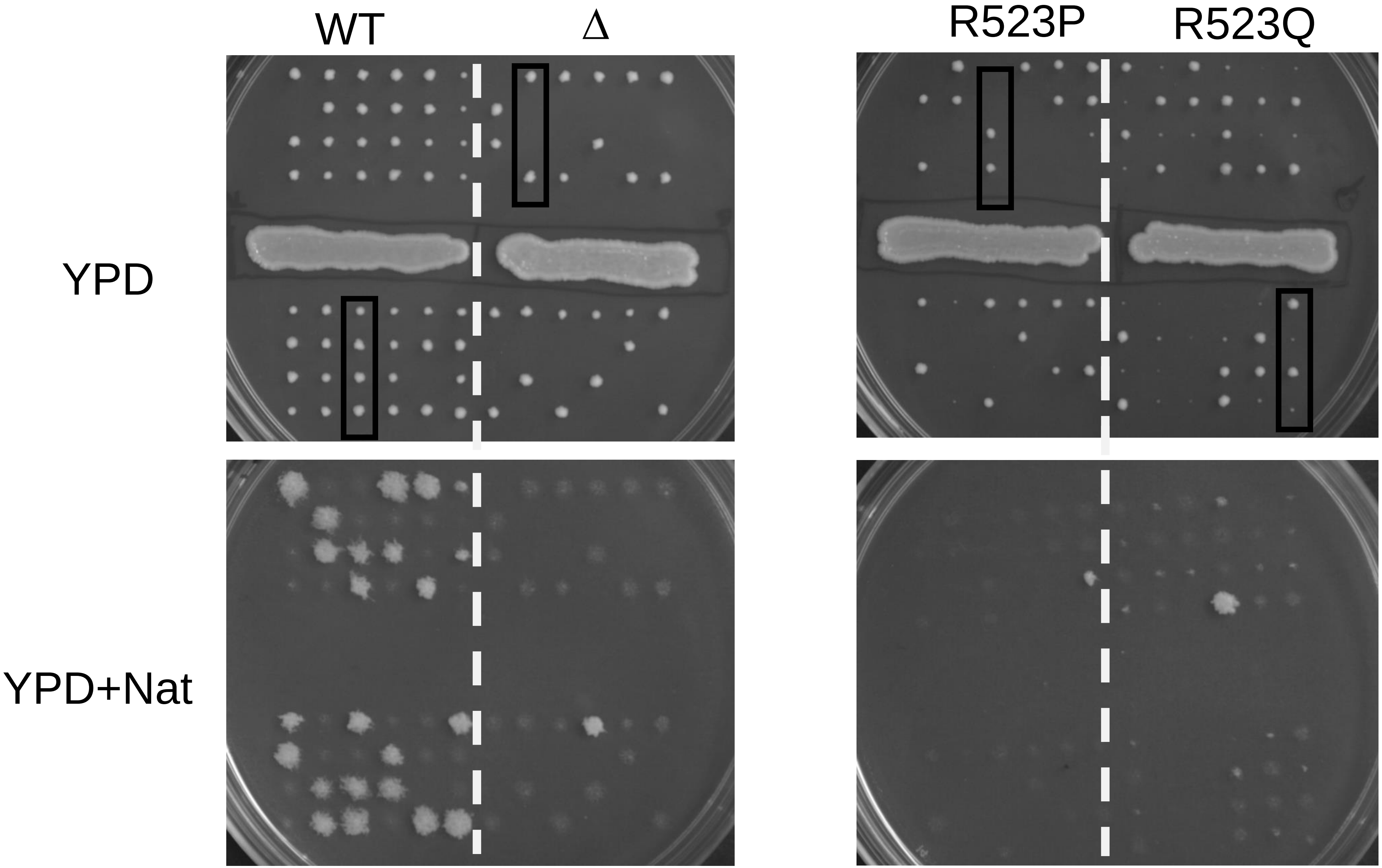

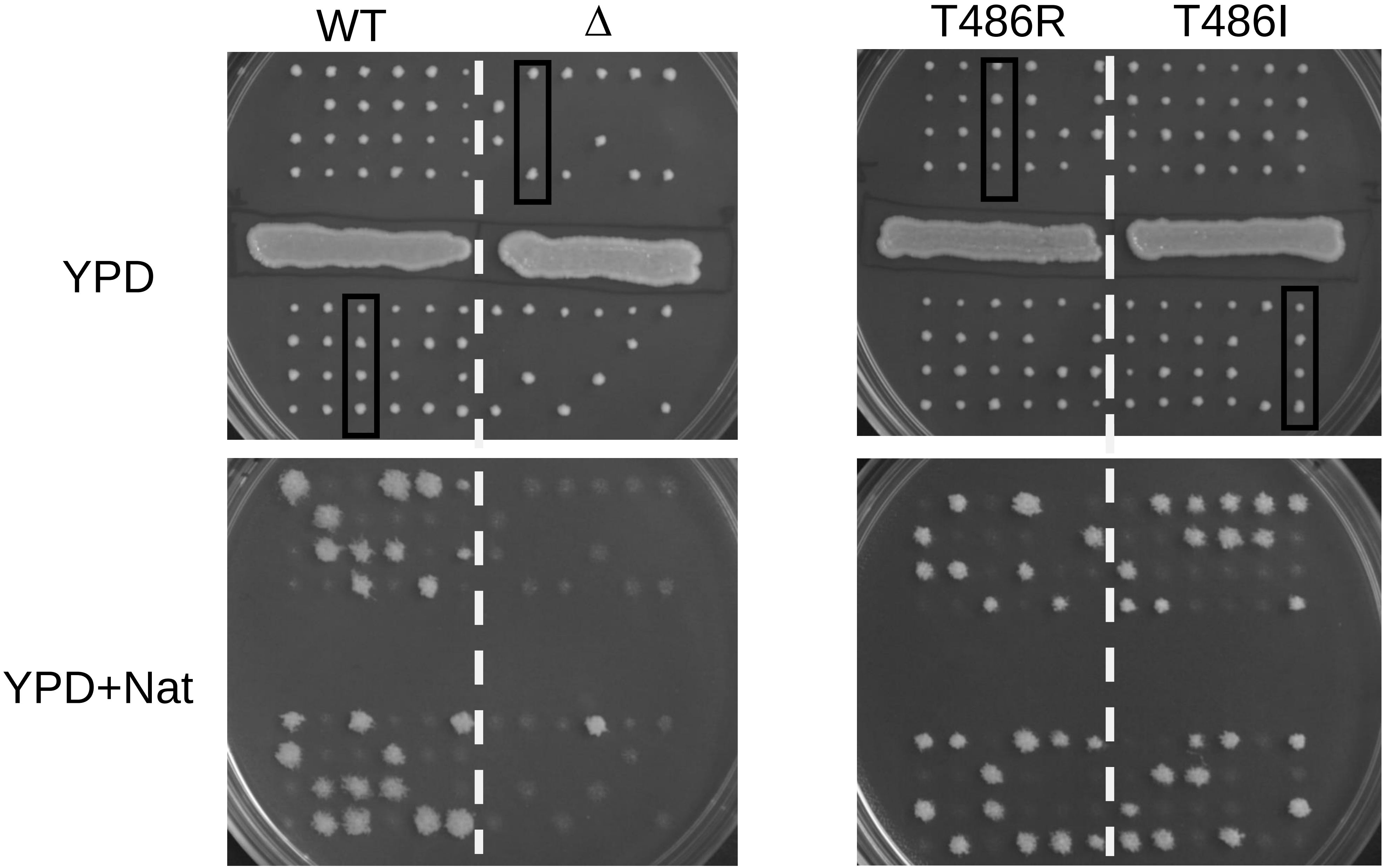

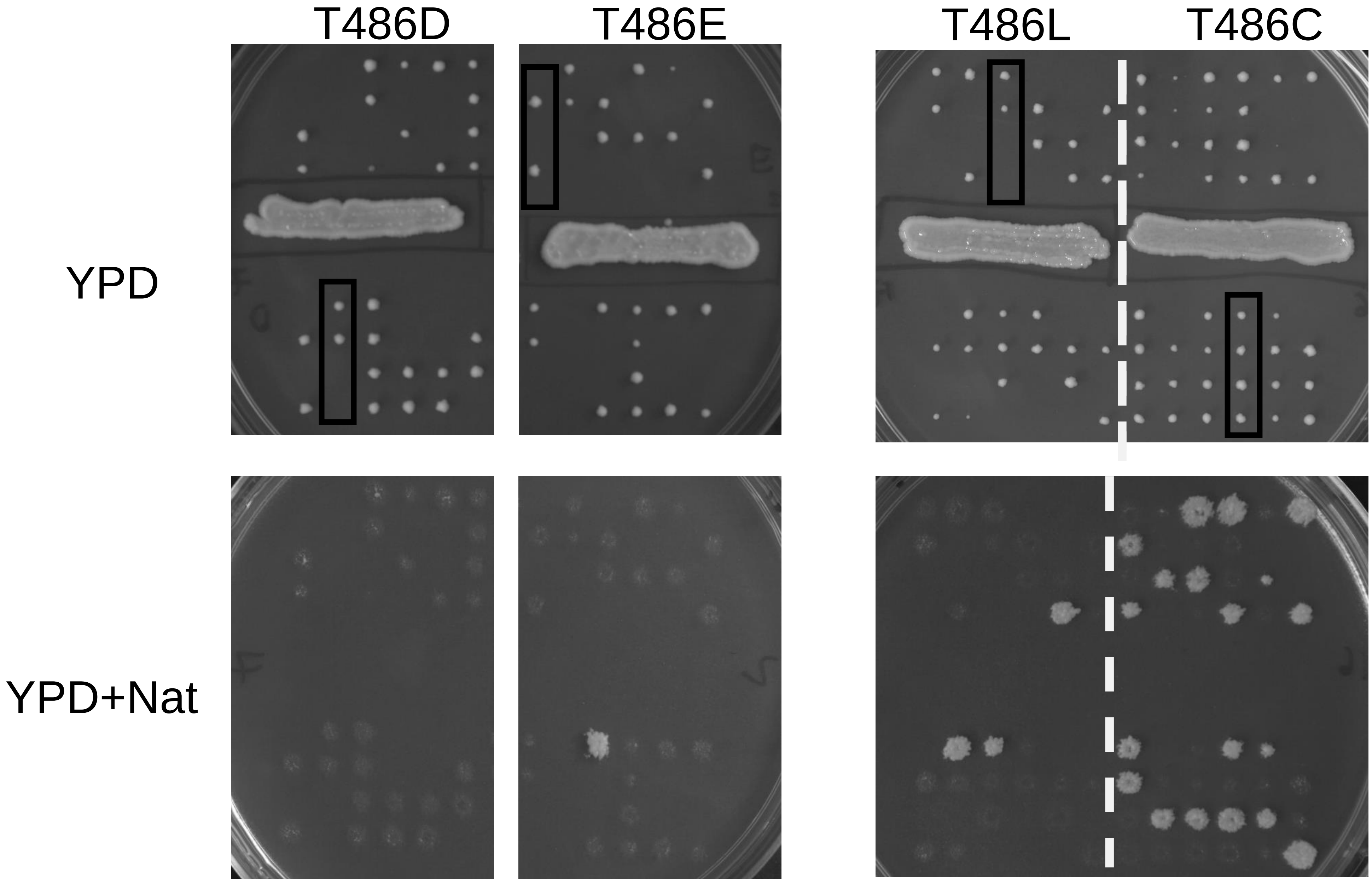

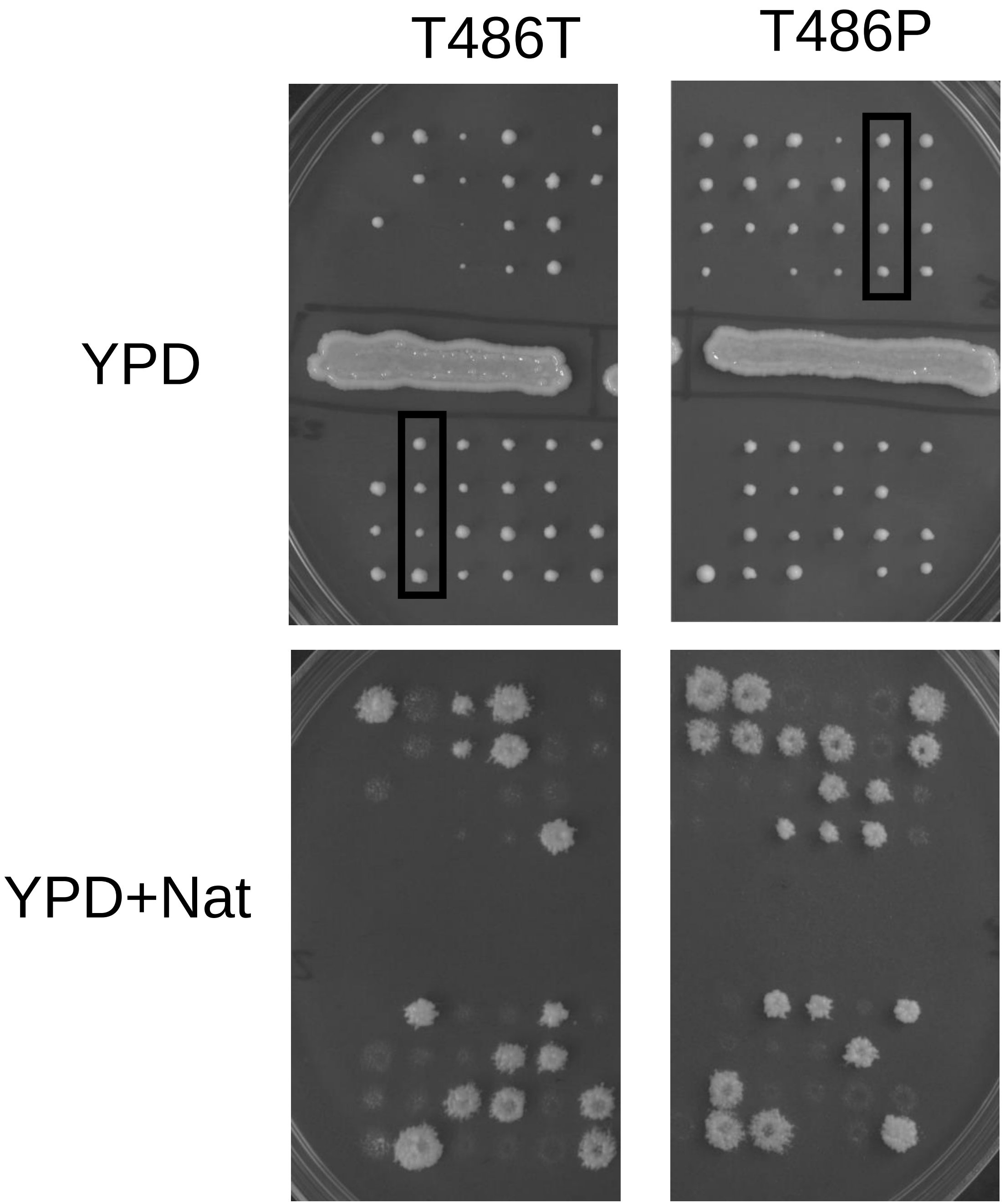
