## Supplementary Figures for "Perturbing proteomes at single residue resolution using base editing"

SUPPLEMENTARY MATERIAL: Supplementary Figures 1-17

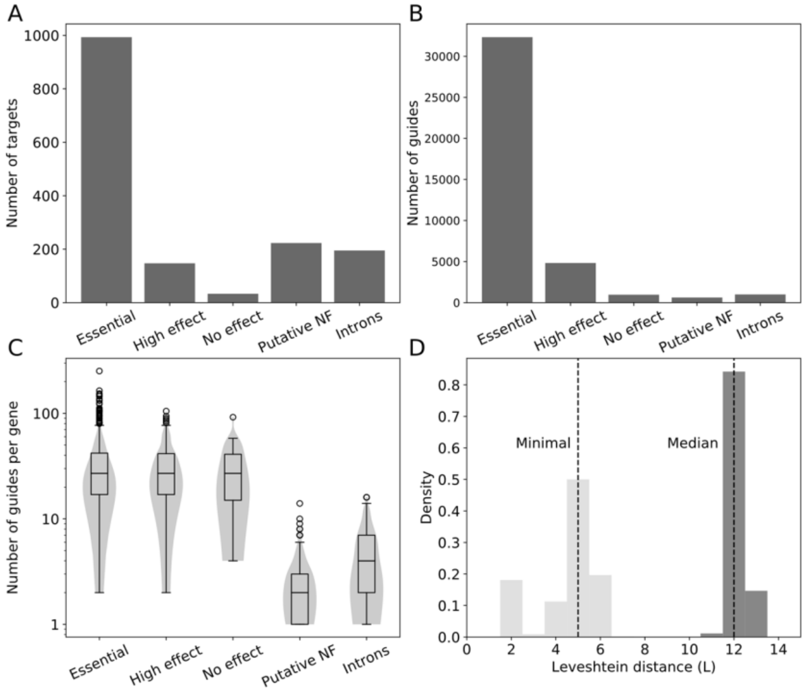


**Supplementary Figure 1: A gRNA library for the systematic mutagenesis of yeast essential genes and other targets of interest. A)** Number of genes targeted by the gRNA library for the different target classes: Essential for essential genes, High effect for genes with large fitness effects when deleted, genes for which No effect on fitness is seen upon deletion, Putative non functional peptides (NF) and Intronic sequences. **B)** Total number of gRNAs targeting genes in the different target classes. Same classes as in A). **C)** Distribution of number gRNAs for each gene targeted in the different classes. **D)** Distribution of minimal (light grey) and median (dark grey) pairwise sequence distance between all gRNA sequences in the library.

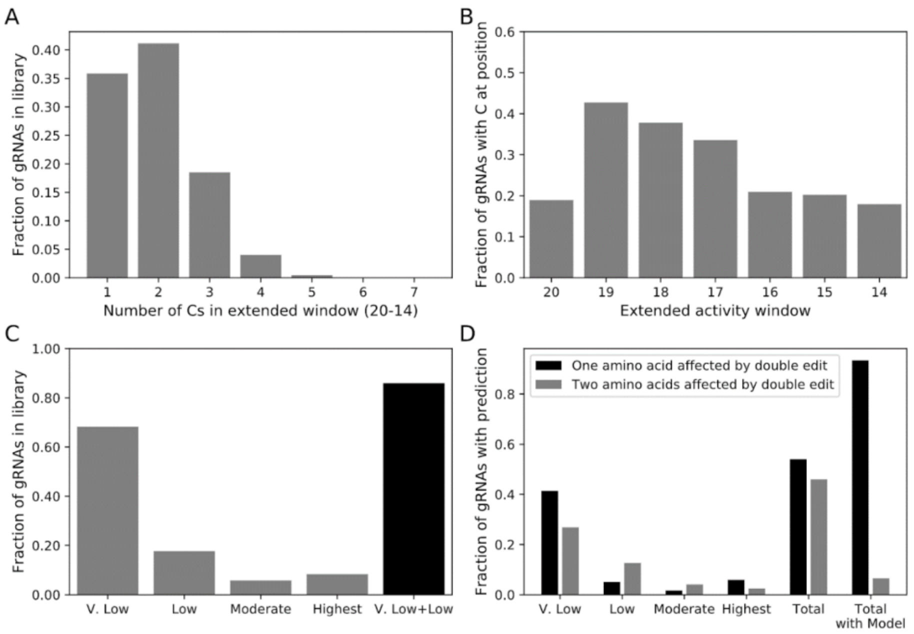


**Supplementary Figure 2 A gRNA library designed to minimize co-editing occurrence and consequences A)** Distribution of the number of editable nucleotides in the extended Target-AID activity window for gRNAs in the library. **B)** Overall occupancy of cytosines in the extended Target-AID activity window (-20 to-14 from the PAM) across the library. **C)** Fraction of gRNAs in different co-editing risk strata based on previous Target-AID data and the deep sequencing data of the present study. The Very Low (V. Low) category represents gRNAs for which only one cytosine is present within the 19 to 16 position range. The Low category represents guides with only one cytosine present within the 19 to 17 position range as well as a cytosine at position 16. The moderate category represents gRNAs with cytosines present at both position 19 and 17, with the possibility of position 16 being a cytosine as well. Finally, the High category includes any guide with cytosines at both position 19 and 18. Over 80% of gRNAs in the library are in either the Very Low or Low co-editing risk categories. **D)** Impact of co-editing on the targeted coding sequence for the different co-editing risk categories. If relative editing rates in the Target-AID activity window are not taken into account, almost half of the gRNAs appear at risk to induce the creation of double mutants at high frequency (Total). If the low rate of editing at the putative co-editing sites of gRNAs in the Very Low and Low risk categories is considered, then over 90% of gRNAs in the library should affect only one amino acid in the target coding sequence even if co-editing occurs.

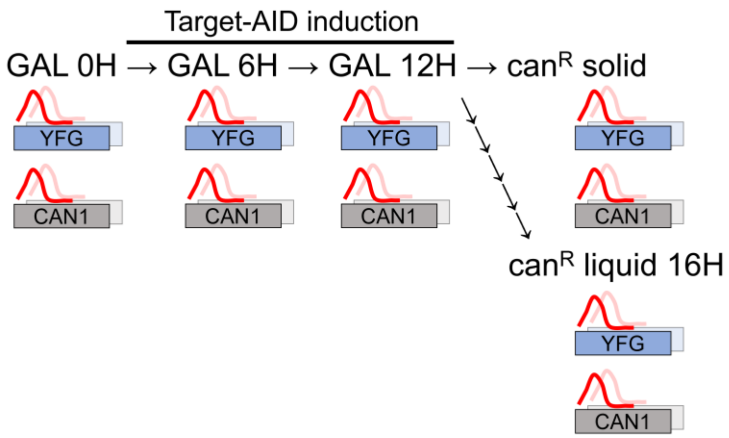


**Supplementary Figure 3 Workflow of the deep sequencing experiment.** After pre-culture rounds in glucose and glycerol synthetic media, cells are placed in galactose containing media, which induces the expression of the Target-AID base editor. After 12 hours, cell dilutions were plated on canavanine containing solid media or put in canavanine containing liquid media for selection of canavanine resistant cells ($can^R$). At each timepoint shown above, cells were sampled, and DNA was extracted. Targeted amplification and sequencing were performed for both the target locus (YFG in the figure) and the co-selection site in CAN1.

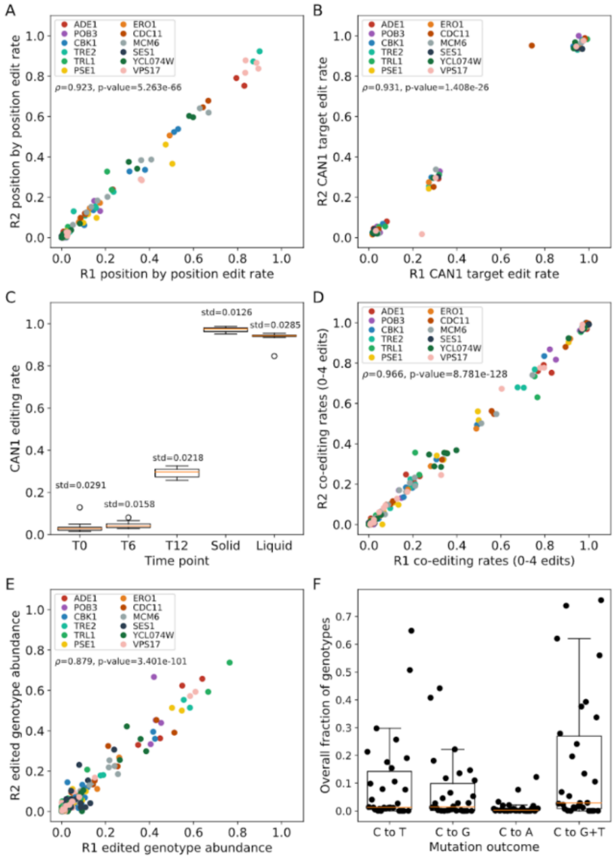


**Supplementary Figure 4 Replication metrics of the deep sequencing experiments A)** Correlation plot for position-wise editing rates observed at all timepoints and editable sites in the experiment. Spearman's rank correlation was calculated using 160 sites/timepoint combinations. **B)** Correlation plot for the CAN1 co-selection target site editing rate for all samples and timepoints in the experiment. Spearman's Rank correlation was calculated with n=58 site/time point combinations. **C)** Distribution of averaged CAN1 editing rates for all samples at the different timepoints (n=12 for all time points except Liquid recovery where n=11). **D)** Correlation plot of co-editing rates (n=1, 2, 3 or 4 edits) for all timepoints and samples. Spearman's rank correlation was calculated using 220 coediting rates/time point combinations. **E)** Correlation plot of base editing outcome relative genotype abundance for all sample and time points. Spearman's rank correlation was calculated using 310 relative genotype abundances/time point combinations. **F)** Occupancy of C toT, C to G and C to A mutations in mutagenesis outcomes weighted by overall site editing rate. The difference between weighted C to T and C to G occupancy is not significant ($W$=162, p=0.15), but the difference between C to T/G and C to A is (C to T vs C to A: $W$=0, p=$1.73x10^{-6}$, C to G vs C to A: $W$=58, p=0.0003).

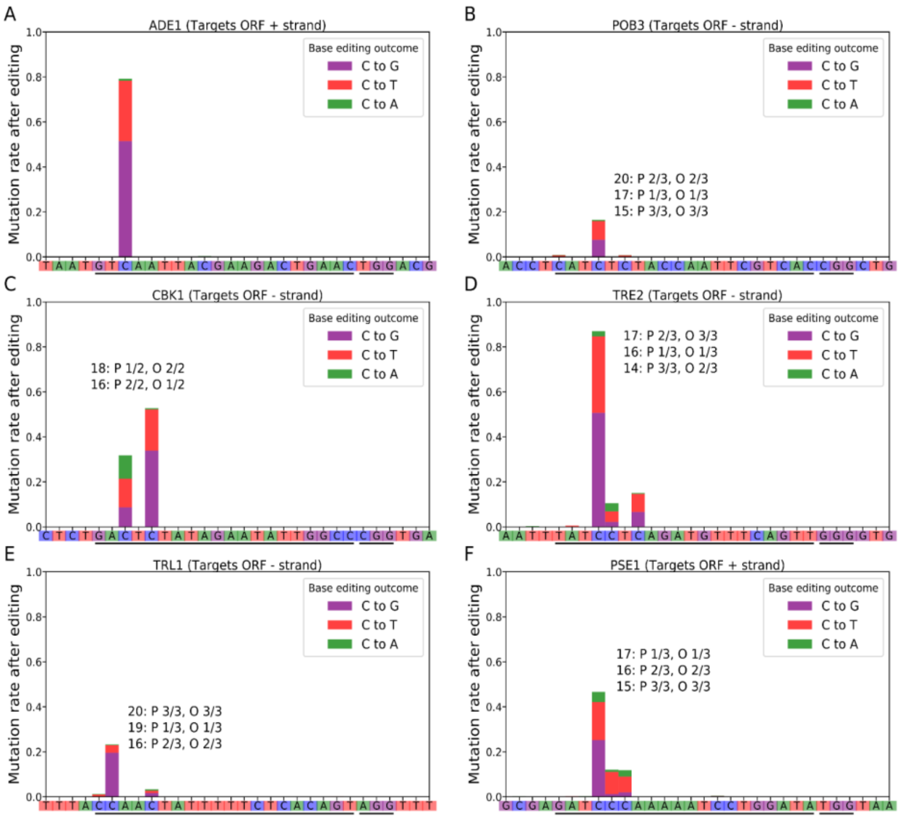


**Supplementary Figure 5 Site specific mutation rates and outcomes of targets 1-6 after co-selection.** The two black lines represent the guide sequence and the PAM. Predicted and observed rankings for editing rates are shown for each gRNA, where P is the predicted ranking and O is the observed ranking. **A)** ADE1 **B)** POB3 **C)** CBK1 **D)** TRE2 **E)** TRL1 **F)** PSE1.

**Commented [P1]:** Changed as part of response to reviewer

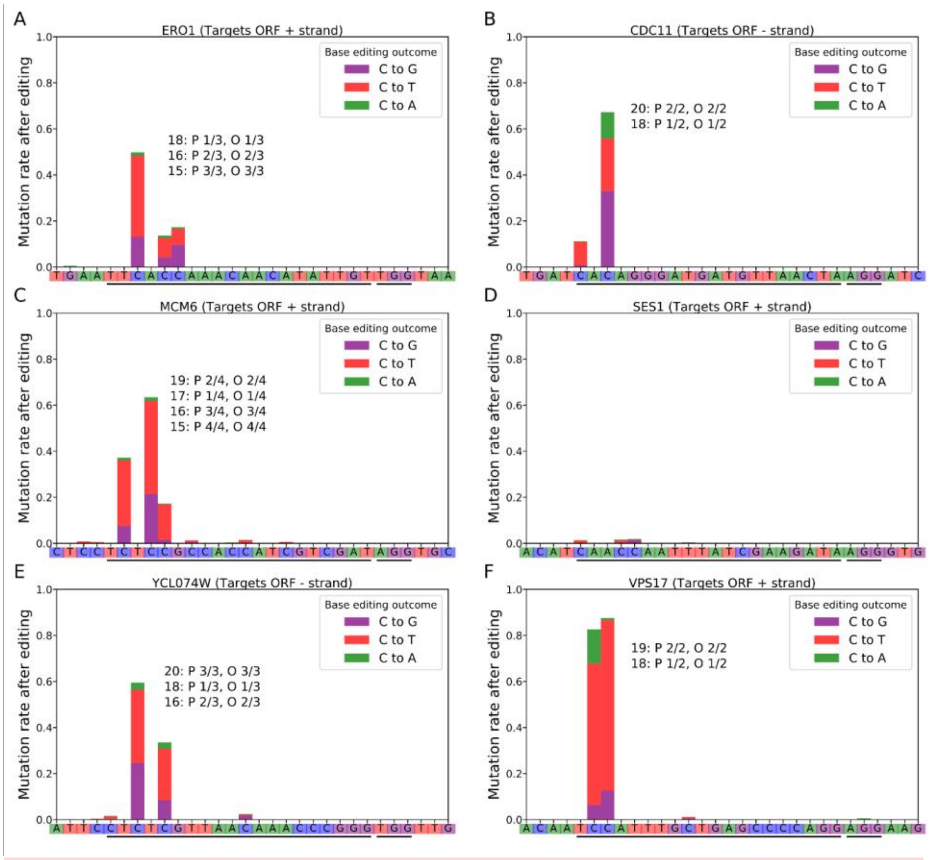


**Supplementary Figure 6 Site specific mutation rates and outcomes of targets 1-12 after co-selection.** The two black lines represent the guide sequence and the PAM. Predicted and observed rankings for editing rates are shown for each gRNA, where P is the predicted ranking and O is the observed ranking. As data from the SES1 target site was not used in the site mutation rank analysis, they are not shown. **A)** ERO1. Because data could not be acquired for the liquid co-selection timepoint, the canavanine media plating data was used instead to generate the figure. **B)** POB3 **C)** CBK1 **D)** TRE2 **E)** TRL1 **F)** PSE1.

**Commented [P2]:** Changed as part of response to reviewer

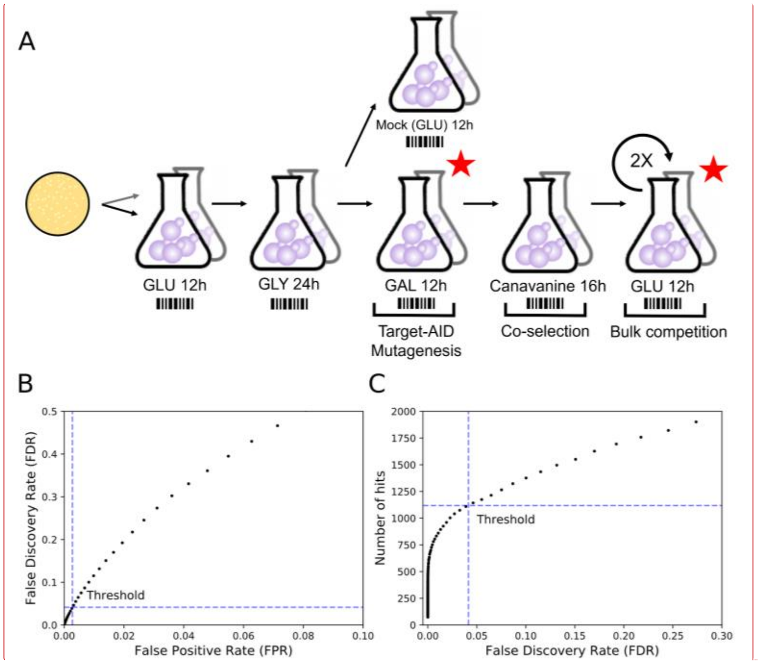


**Supplementary Figure 7: Experimental workflow for Target-AID mutagenesis and co-selection. A)** The mutagenesis method closely follows the base editing protocol previously described[23]. After a pooled transformation, cells were scraped from the solid media plates and splitted into two replicates for pre-cultures. After each step of the protocol, plasmid DNA was extracted from a cell sample and used to amplify and sequence the plasmid inserts corresponding to the gRNAs. The red stars indicate time points used for fitness effects analysis: read counts after galactose induction were used as T0 and were compared with read counts after two rounds of competition. The mock induction steps mimic the induction conditions but galactose in the media is replaced by glucose. This prevents the editing enzyme from being expressed because glucose represses the GAL pathway. After canavanine co-selection, cells go through two competition rounds in synthetic media where selective pressure for the Target-AID bearing plasmid is lost. The entire experiment was completed within less than 25 generations after galactose induction, limiting the impact of compensatory and spontaneous mutations. **B)** Calculated False Discovery Rates (FDR) as a function of the False Positive Rate (FPR) Threshold set to select gRNAs with significant drops in abundance based on the Glucose to Mock reference distribution. **C)** Number of gRNAs with significant drops in abundance (GNEs) as a function of the FDR threshold set.

**Commented [P3]:** Changed as part of review

**Commented [P4]:** Added as part of review

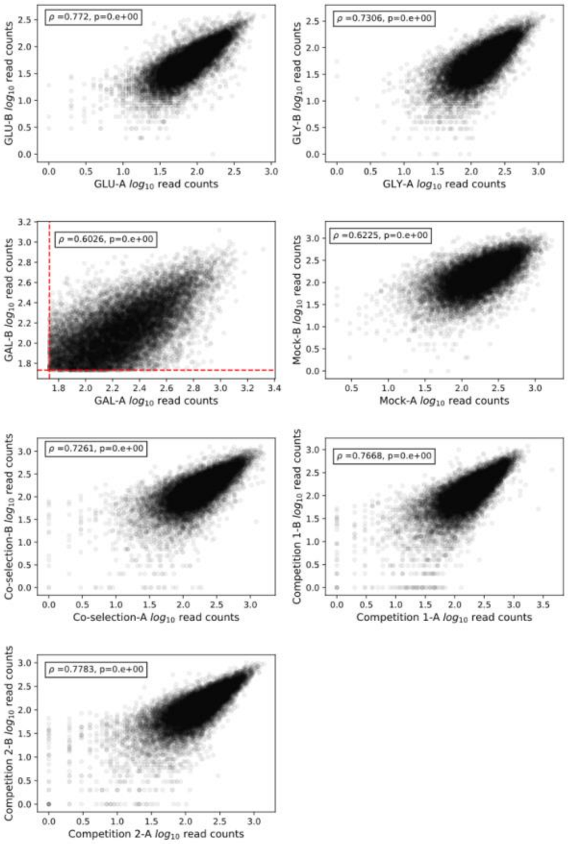


**Supplementary Figure 8: Read abundance rank order is strongly correlated between replicates.** For each time point, Spearman rank correlation of gRNA $log_{10}$ read abundance after basic filtering is shown. The minimal read count after galactose induction, which served as the principal filtering criteria, is shown on the galactose subpanel.

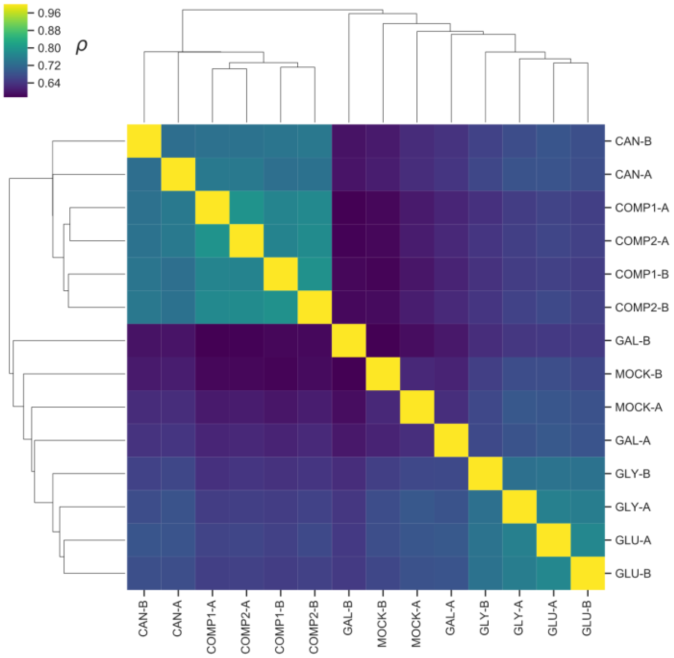


**Supplementary Figure 9: Barcode abundance correlation clusters different experimental steps of the screen.** Pairwise Spearman rank correlation of barcode counts was used to cluster the libraries obtained at the different time points described in Figure S2. The lower level of correlation between the galactose induction and mock induction timepoints compared to other associated steps could reflect higher stochasticity in growth caused by cell to cell variation in the metabolic switch from glycerol to sugars as the main carbon source as well as editing in the case of the galactose timepoint.

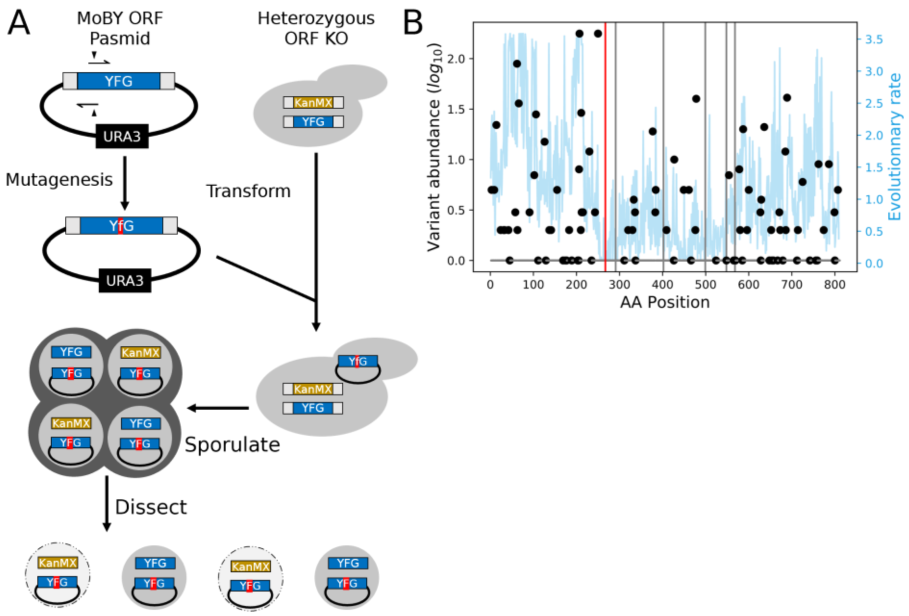


**Supplementary Figure 10 Plasmid-based confirmation workflow by complementation test and evolutionary information on *GLN4*. A)** Detailed protocols for the different steps are presented in the methods. First, site directed mutagenesis is used to introduce the mutation of interest (shown in red) in the MoBY collection plasmid of the targeted gene (YFG). This vector is then transformed into the heterozygous collection deletion strain (BY4743, *MAT*a/α *his3Δ1/his3Δ1 leu2Δ0/leu2Δ0 LYS2/lys2Δ0 met15Δ0/MET15 ura3Δ0/ura3Δ0*) of the gene of interest. The transformants are sporulated and their tetrads are dissected. If the mutated allele carried by the plasmid cannot complement the gene deletion, then only the two progenies bearing the wild-type copies will be viable. **B)** Protein variant frequency among 1000 yeast isolates (black dots) and residue evolutionary rate across species (blue line) for *GLN4*. The target site for the most deleterious GNE is highlighted by a red line and other GNE target sites are shown as grey lines.

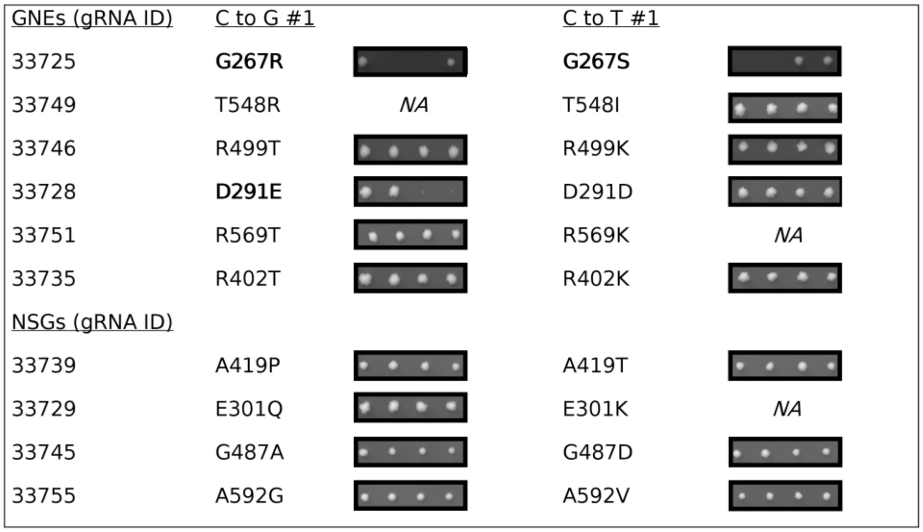


**Supplementary Figure 11 Validation studies of GNEs and NSGs targeting *GLN4.*** Tetrad dissection patterns of the most probable mutagenesis outcomes for GNEs targeting *GLN4* as well as 4 NSGs targeting amino acids close to the GNE target sites. NA denote sites which were not considered further out after unsuccessful directed mutagenesis.

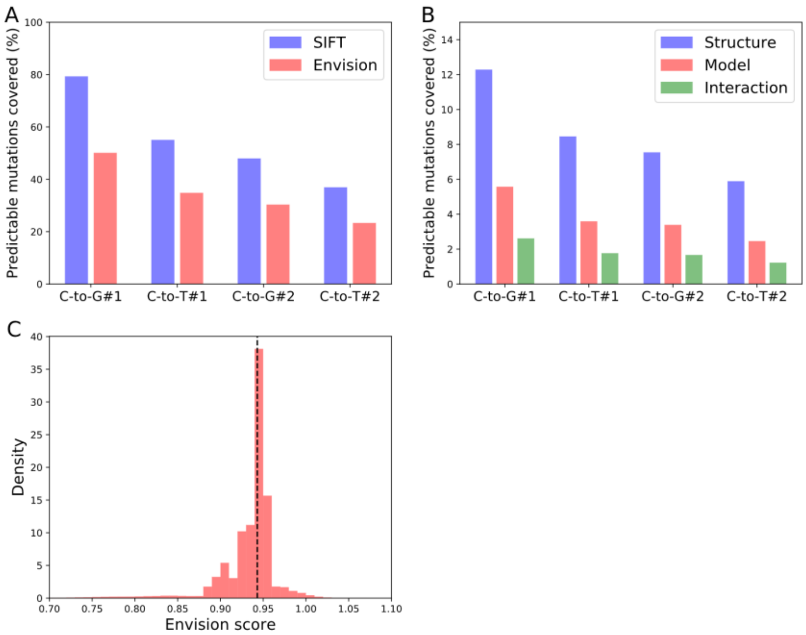


**Supplementary Figure 12 gRNA predicted mutation coverage for Mutfunc and Envision data.** Mutfunc integrates both the SIFT prediction scores and FoldX[68], ΔΔG predictions for solved protein structures, homology models, and protein-protein interaction interfaces. gRNAs which do not generate missense mutations were included in the calculations. **A)** Coverage for the SIFT and Envision variant effect predictors for the four most probable single mutants created by gRNAs detected in the experiment. **B)** Coverage for ΔΔG predictions for solved protein structures, homology models, and protein-protein interaction interfaces for the four most probable single mutants created by gRNAs detected in the experiment. **C)** Distribution of Envision scores across all sites in the database for all proteins targeted by the set of gRNAs detected in the screen (n=7,556,573). The median score is shown as a dotted black line.

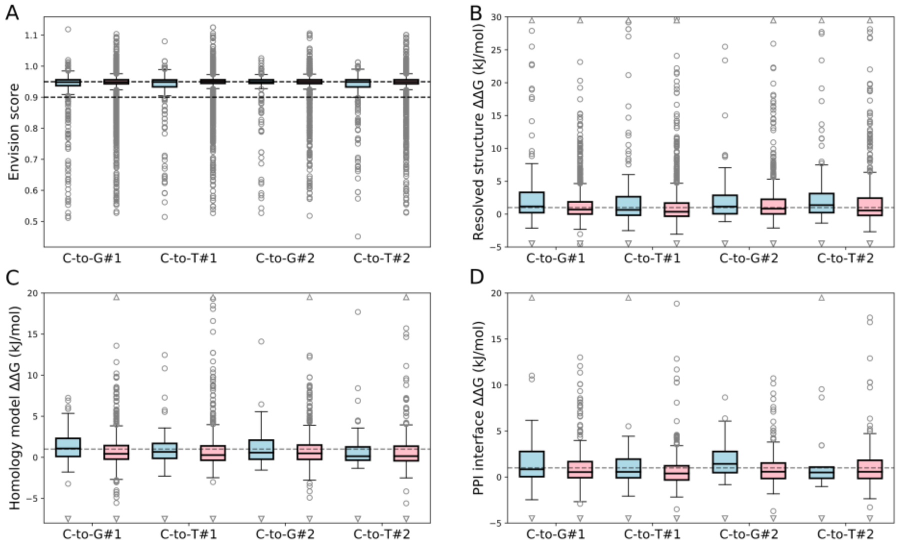


**Supplementary Figure 13 GNE and non-significant gRNA effect prediction distributions. A)** Envision score distributions for the four most probable mutations induced by GNEs (blue) and NSGs (red) (n= 547, 7850, 424, 5410, 294, 3615, 390, 4694). Welch's t-test p-values for comparisons: $5.00 \times 10^{-6}$, 0.002, 0.007, $7.75 \times 10^{-5}$. **B)** Predicted folding energy variation (ΔΔG) of GNE and NSG induced protein mutants compared to the wild-type structure based on resolved protein structure (n= 168, 1890, 128, 1289, 113, 1151, 88, 899). Welch's t-test p-values for comparisons: 0.0001, 0.006, 0.148, 0.007. **C)** Predicted folding energy variation (ΔΔG) of GNE and NSG induced protein mutants compared to the wild-type structure based on homology models of protein structure (n= 88, 846, 56, 546, 59, 509, 39, 373). Welch's t-test p-values for comparisons: 0.016, 0.441, 0.195, 0.689. **D)** Binding energy variation (ΔΔG) of GNE and NSG induced mutant protein-protein interfaces compared to the wild-type based on a resolved structure of the interface (n= 41, 397, 30, 266, 27, 252, 21, 184). Welch's t-test p-values for comparisons: 0.285, 0.303, 0.033, 0.95.

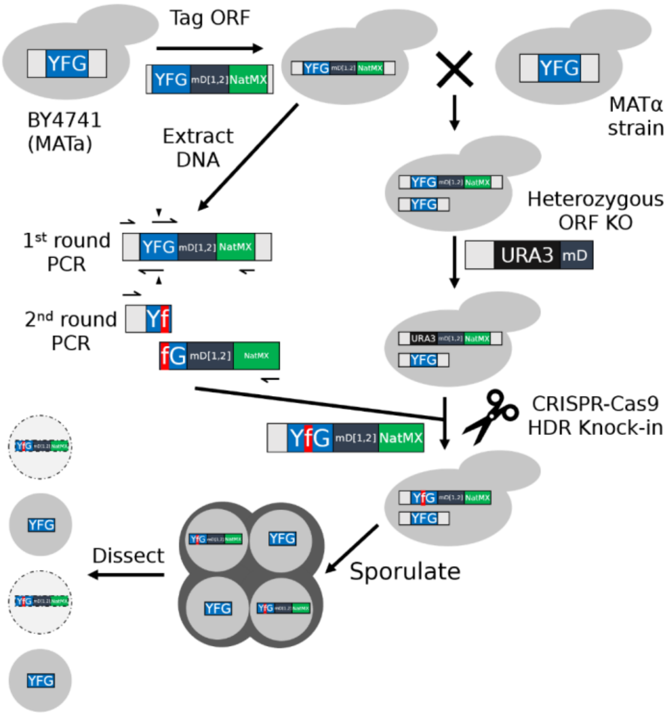


**Supplementary Figure 14 Fitness affecting variant by CRISPR knock in validation workflow.** Detailed protocols for the different steps are presented in the methods. Starting from the wild-type laboratory strain BY4741, the gene of interest (*YFG*, blue) is first tagged with a modified version of the DHFR F[1,2] cassette (dark gray and green). The tagged strain is then crossed with a MAT*α* strain (Y8205) to create a heterozygous diploid. A *URA3* deletion cassette (black) that recombines with the *YFG* upstream sequence and the start of the mDHFR fragment is then used to generate a heterozygous KO strain. In parallel, genomic DNA is extracted from the tagged haploid strain. This DNA is then used as a template to amplify two fragments of *YFG* bearing the mutation of interest (shown in red) using a set of overhanging primers. The two fragments are then combined by fusion PCR to obtain the donor DNA used in the next step. Using a modified Cas9 vector[60] that expresses a gRNA targeting the *URA3* cassette, the mutated allele is introduced at the KO locus to create a heterozygous mutant strain. The diploid cells can then be sporulated, and tetrad dissection allows observation of any phenotype linked with the mutation of interest.

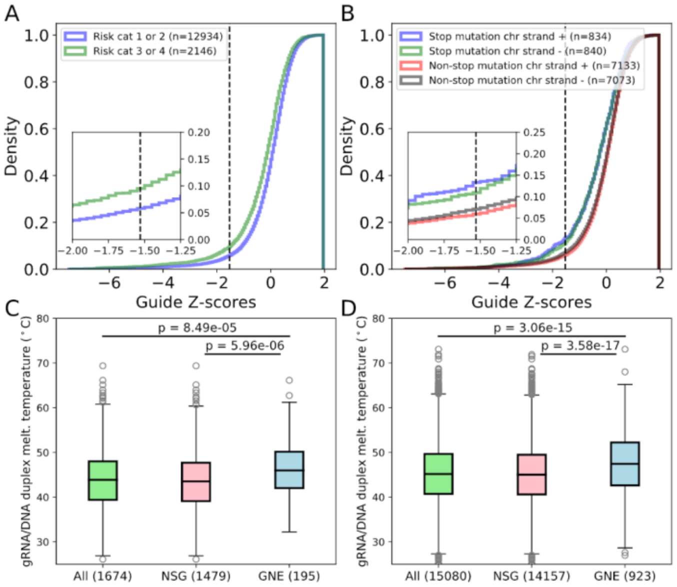


**Supplementary Figure 15 Other properties of SGG and NSG GNEs. A)** Cumulative z-score density for gRNAs that do not generate stop codons depending on the co-editing risk category. A higher rate of GNE is observed for gRNAs which can lead to the editing of multiple nucleotides (Two-sample Kolmogorov-Smirnov test, p=2.32x10-24). The significance threshold is shown as a black dotted line. **B)** gRNA z-score cumulative density for both SGGs and non-SGGs grouped by the chromosomal strand they target. In SGGs, the target strand does not impact z-score distributions (Two-sample Kolmogorov Smirnov test, p=0.717) and GNE proportions (Fisher's exact test, p=0.149). For non-SGGs, the chromosomal strand has a small influence on z-score distributions (Two-sample Kolmogorov Smirnov test, p=0.04) and GNE proportions (Fisher's exact test, p=0.002) **C)** Distributions of modeled RNA/DNA duplex melting temperature for all SGGs, the NSG subset, and the GNE subset. P-values were calculated using the two-sample Kolmogorov Smirnov test. **D)** Distributions of modeled RNA/DNA duplex melting temperature for all non-SGGs, the NSG subset, and the GNEs subset. P-values were calculated using the two-sample Kolmogorov Smirnov test.

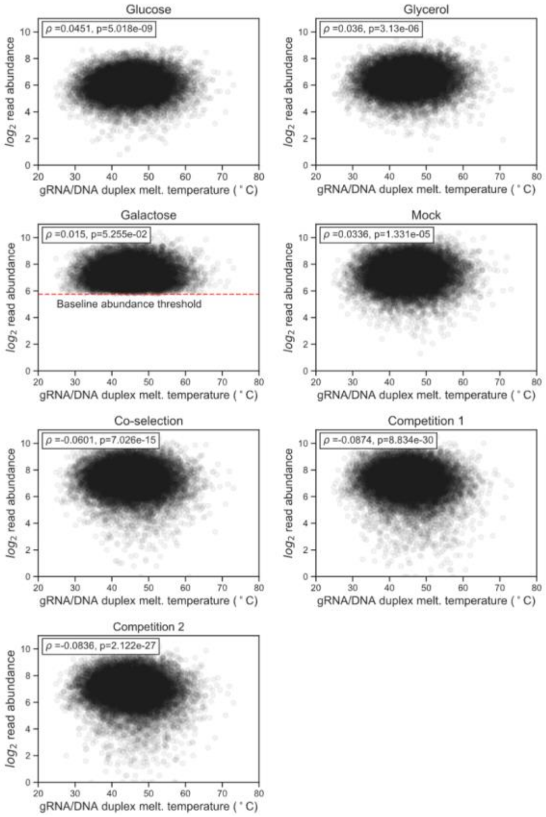


**Supplementary Figure 16 gRNA/DNA duplex melting temperature is not linked to systematic sequencing biases.** Spearman rank correlation between replicate averaged read counts and predicted gRNA/DNA duplex melting temperature is shown across timepoints. The minimal read count after galactose induction, which served as a filtering criterion, is shown on the galactose subpanels. gRNAs for which no reads were detected in one of the time points were included when computing the correlation but are not shown on the graphs because of log scaling.

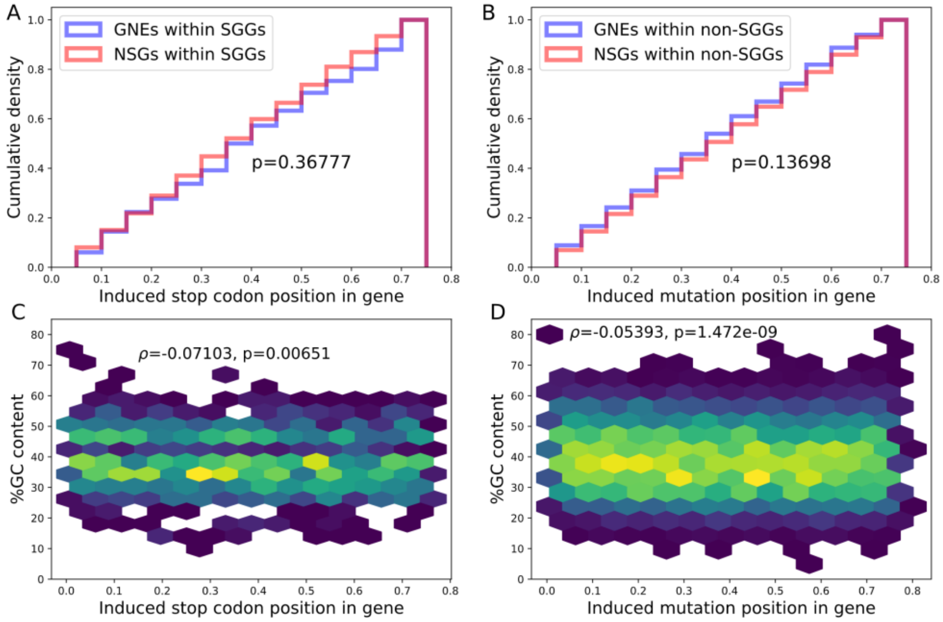


**Supplementary Figure 17 GNE density is independent of target nucleotide position bias. A)** In Stop codons generating gRNAs (SGGs), gRNAs with significant negative fitness (GNEs) and gRNAs with no significant effects (NSGs) target sites are evenly distributed across the target genes, and GNEs do not show any bias (Two-sample Kolmogorov-Smirnov). **B)** Non-SGG GNEs do not show any positional bias. **C)** A significant but small negative correlation is observed between gRNA target relative position and GC content of SGGs (Spearman's rank correlation). The very small observed effect coupled with the absence of position bias suggests that relative target position bias does not drive the link between GC content and gRNA efficiency. **D)** Similarly, a small but significant but small negative correlation is also observed between gRNA relative position and GC content for non-SGGs (Spearman's rank correlation).
